# Defining the mechanisms and properties of post-transcriptional regulatory disordered regions by high-throughput functional profiling

**DOI:** 10.1101/2024.02.01.578453

**Authors:** Joseph H. Lobel, Nicholas T. Ingolia

## Abstract

Disordered regions within RNA binding proteins are required to control mRNA decay and protein synthesis. To understand how these disordered regions modulate gene expression, we surveyed regulatory activity across the entire disordered proteome using a high-throughput functional assay. We identified hundreds of regulatory sequences within intrinsically disordered regions and demonstrate how these elements cooperate with core mRNA decay machinery to promote transcript turnover. Coupling high-throughput functional profiling with mutational scanning revealed diverse molecular features, ranging from defined motifs to overall sequence composition, underlying the regulatory effects of disordered peptides. Machine learning analysis implicated aromatic residues in particular contexts as critical determinants of repressor activity, consistent with their roles in forming protein-protein interactions with downstream effectors. Our results define the molecular principles and biochemical mechanisms that govern post-transcriptional gene regulation by disordered regions and exemplify the encoding of diverse yet specific functions in the absence of well-defined structure.

## Introduction

Over one third of eukaryotic proteins contain intrinsically disordered regions that vary in length, composition, molecular properties, and biological function^1,2^. Although they lack well defined structures, disordered regions nonetheless potently and specifically regulate diverse cellular processes^1,2^. Biochemical studies have revealed that functional disordered regions achieve their regulatory effects through varied molecular interactions with binding partners, ranging from short linear motifs, as seen in proline rich stretches recognized by SH3-domains, to fuzzy binding using overall sequence composition, exemplified by transcription factor activation domains^2–6^. These functional disordered regions are especially prominent within RNA binding proteins (RBPs), where they engage binding partners to tune the stability and translation of target mRNAs^7,8^. While the requirement of disordered regions within RBPs for regulatory activity is well appreciated, the biochemical properties and molecular mechanisms driving their function remains poorly understood. Determining the pathways and sequence features used by these regulatory disordered regions is essential for understanding how cells control mRNA stability and translation.

Comprehensive identification of RNA-binding proteins has revealed thousands of potential post-transcriptional regulators^9^. These proteins contain a combination of structured domains and intrinsically disordered regions that cooperate to execute specific regulatory outcomes^10^. While the folded domains of RBPs bind target sequences within transcripts, the unstructured regions can recruit effectors, such as translation or mRNA decay factors, to the mRNA through protein-protein interactions^8^. Structured domains and disordered regions may thus constitute separable functions of the protein that collectively regulate the post-transcriptional fate of an mRNA, much like transcription factors use folded DNA binding domains to interact with genomic elements and disordered regions to recruit coactivators^11^. Understanding the contributions of both folded domains and disordered regions is required for a complete framework of how post-transcriptional regulators sculpt mRNA dynamics.

While classic investigations of full-length RNA-binding proteins or isolated structured domains have been invaluable for discovering the specific mRNAs bound by RBPs, the contribution of the ubiquitous disordered regions has remained less clear^12–16^. Compared to globular regions, functional annotation of disordered elements remains difficult because these elements are often poorly conserved and vary considerably in length and composition across evolution^17^. Consequently, these studies typically focus on select motifs within disordered regions demonstrating how these sequences engage downstream machinery to influence the translation and stability of target messages. Disrupting these motifs often cripples regulatory function of the full-length protein; reciprocally, tethering the isolated motif to a reporter mRNA can be sufficient for activity, highlighting the key contributions of these disordered regions to post-transcriptional regulation^18–20^. More broadly, characterization of functional disordered regions across diverse biochemical pathways have revealed that these elements employ a range of molecular grammars—from defined short linear motifs that co-fold through specific interactions to overall sequence composition that engage in fuzzy binding—to mediate their regulatory effects^1,2^. While individual examples are informative, the full scope of how disordered regions drive post-transcriptional regulation remains elusive.

In this study, we functionally profile the post-transcriptional regulatory activity of ∼45,000 disordered peptides spanning the disordered proteome of budding yeast. Our survey leverages a fluorescence-based tethered function assay that enables pooled, high-throughput activity measurements in yeast. We identified hundreds of elements that regulate mRNA stability and translation and reconstruct functionally important disordered regions within post-transcriptional regulators. By profiling regulatory activity in genetic backgrounds lacking key mRNA decay nucleases, we determine the molecular pathways underlying the activity of many repressive disordered peptides. We then precisely define critical residues within these disordered regions by mutational scanning and characterize the diverse sequence features employed by regulators, which range from local motifs to overall amino acid composition. Computational and machine learning analysis of these large-scale functional data reveals a major contribution by aromatic residues to the regulatory function of these disordered regions and highlights important residues based on sequence context. This comprehensive analysis enables us to define the molecular rules and mechanisms used by disordered regions to control mRNA stability and protein synthesis.

## Results

### High-throughput measurement of post-transcriptional regulation by disordered regions

To broadly survey how the disordered proteome modulates mRNA stability and translation and systematically explore sequence-function relationships, we developed a high-throughput, fluorescence-based approach to measure post-transcriptional regulatory activity^12^. Our method builds upon classic RNA tethered-function assays that use a heterologous RNA-protein interaction to recruit a query protein to a reporter transcript and examine how the tethered construct affects mRNA decay and protein synthesis^12,13,21,22^. We targeted query proteins to a reporter transcript that encodes a yellow fluorescent protein (YFP) by fusing these proteins to the 22 amino acid λN phage peptide, which recognizes boxB RNA hairpin motifs that we embed in the 3ʹ UTR of the YFP mRNA^23^. Tethered fusion proteins that promote mRNA decay or repress translation should lower the YFP output. We normalize YFP fluorescence against a non-targeted red fluorescent mScarlet reporter to correct for cell size and non-specific effects, enabling quantitative ratiometric detection of repressor activity by flow cytometry (**Figure 1A)**^24^. Notably, we designed the reporter mRNA to maximize the dynamic range for YFP signal loss and increase our sensitivity for detecting post-transcriptional repressors by constructing our reporter with optimized codons and stabilizing UTRs^25,26^. We engineered a yeast strain expressing the YFP-boxB reporter mRNA and the mScarlet normalizer and tested our fluorescence-based tethering assay by expressing a repressive, disordered fragment of the nonsense-mediated decay factor, Ebs1, fused to the λN peptide^27^. We observed a ∼5 fold decrease in YFP/RFP ratio relative to λN alone or fused with a non-repressive disordered control, verifying that our system captures the post-transcriptional activity of established regulators **(Figure 1B and S1A-C)**. Thus, this fluorescence-based tethering system couples post-transcriptional regulation to a quantitative measurement of repressor activity.

**Figure 1:**
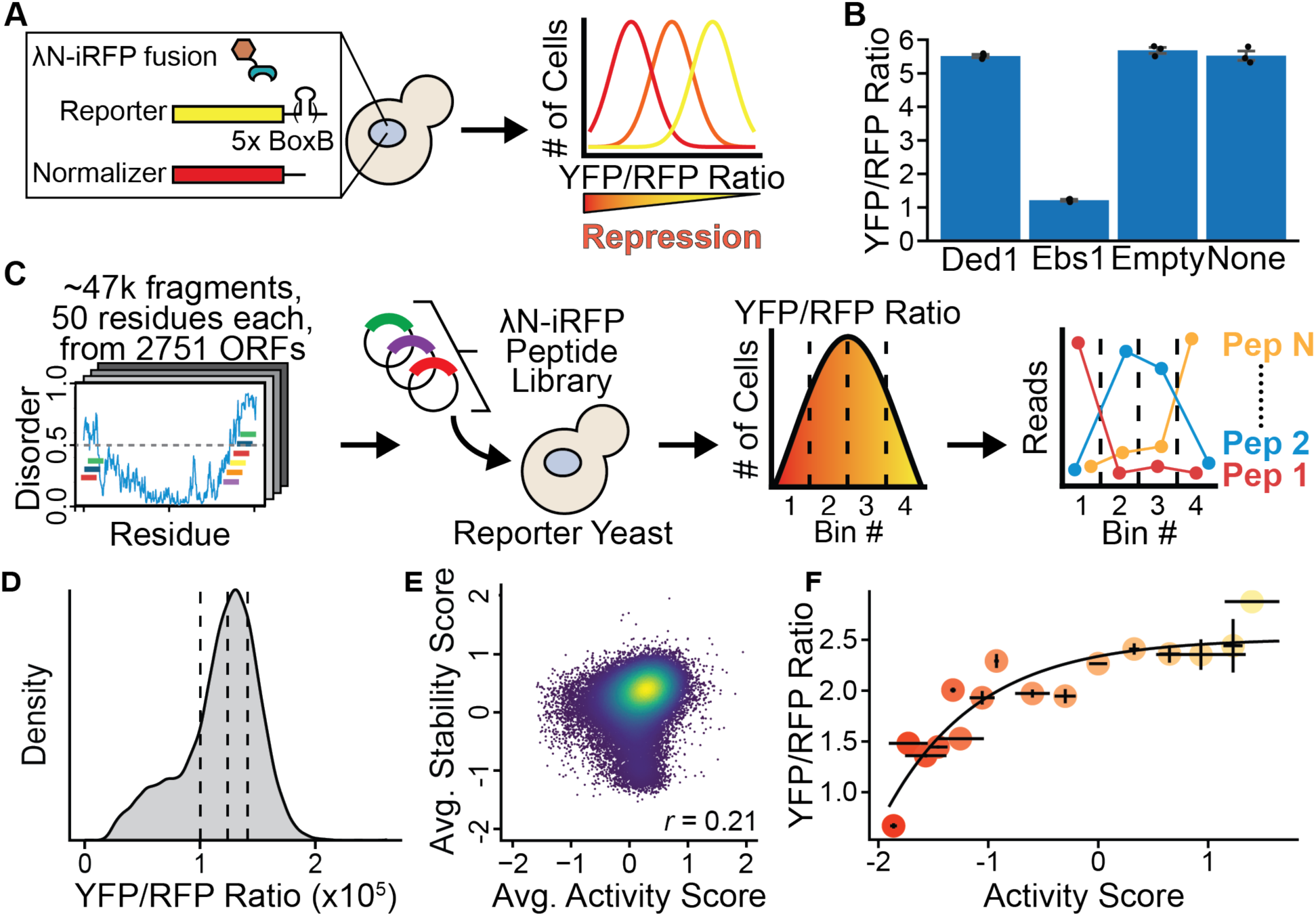
High-throughput functional profiling of post-transcriptional regulatory activity in disordered regions. **(A)** Schematic of fluorescence-based tethering assay. A YFP reporter mRNA contains 5x BoxB hairpins in its 3′ UTR. Query proteins are fused to the λN tethering protein. **(B)** Fluorescence ratio changes of tethered fragments or no tethering construct. For Ded1, residues 1-108 were used. For Ebs1, residues 691-876 were used. Individual mean values and standard deviation are displayed (n=3). **(C)** Schematic of disordered fragment library design and Sort-seq workflow. **(D)** Distribution of YFP/RFP ratio of reporter yeast expressing the full disordered sequence library. Vertical lines demarcate equal population bins used in flow sorting. **(E)** Comparison between average activity and stability scores, with the Pearson correlation coefficient displayed in panel legend (n=2). **(F)** Tethering activity of individually selected fragments compared to activity score by high-throughput analysis with standard deviation (n=2). Correlation was fit with exponential rise to max. Fragments tested, from left to right: Pho92(17-66), Edc1(126-175), Rat1(919-968), Puf4(71-120), Caf120(915-964), Ebs1(825-874), Pbp1(614-663), Spc110(260-309), Sec31(1043-1092), Rad14(101-150), Lam4(351-400), Sac3(689-738), Pry1(113-162), Tif4631(191-240), Pim1(848-897), Gcd11(11-60).

To realize the full potential of fluorescence-based activity measurements, we designed a highly complex library of tethering constructs that spanned all disordered regions in the budding yeast proteome. We chose to screen the entire disordered proteome, as this would provide a comprehensive catalog of potential regulatory regions unbiased by known RNA-binding activity. To develop this library, we predicted the disordered propensity of every residue in the yeast proteome and divided these regions into overlapping 50 amino acid windows. This tiling approach offers precise identification of functional subregions, while testing individual fragments large enough to contain motifs or locally important physicochemical features, and yielded a diverse library of ∼47,000 fragments from 2751 ORFs (∼50% of the yeast proteome) to evaluate for regulatory activity **(Figure 1C)**. We suspected that some fragments might express poorly and therefore show unchanged YFP/RFP ratios because the peptide was unstable, conferring no information about its activity^22^. To distinguish between truly inactive and unstable peptides, we appended a monomeric emiRFP670 to the λN peptide to measure its expression level in a third fluorescence channel **(Figure S1D)**^28^.

We then measured the regulatory activities of this comprehensive disordered peptide library in parallel, by flow sorting yeast expressing tethered peptide constructs and characterizing the sorted cells by high-throughput sequencing (Sort-seq) ^3,29,30^. To perform this Sort-seq analysis, we transformed the yeast strain expressing our reporters with the library of disordered tethering constructs and then grew these transformants at constant density for 16 hours before sorting into four equally sized bins based on the YFP/RFP ratio^31^. The fluorescence ratio distribution showed a clear sub-population with reduced YFP/RFP ratios, indicating the presence of many post-transcriptional repressors within our library **(Figure 1D)**. We quantified the constructs in each bin by deep sequencing and computed an activity score for each fragment based on its read distribution across the bins **(Figure 1C)**. Strongly repressive fragments had a higher portion of reads in bins with lower YFP/RFP ratio, while inactive constructs are found in bins with a higher ratio **(Figure S1E)**. In a complementary approach, we also sorted yeast based on iRFP/RFP ratio to obtain a stability score for each tethering construct **(Figure S1F)**. There was high correlation between biological replicates for both the activity and stability scores, indicating the technical robustness of this assay **(Figures S1G-H)**. Importantly, fragments with low stability scores had near-zero activity, demonstrating how the iRFP measurements enable us to distinguish peptides that lack repressive activity due poor expression **(Figure 1E)**. Overall, this approach captured activity and stability scores for 43,820 tethering constructs to provide a proteome-wide assessment of disordered regions with post-transcriptional regulatory activity.

We next determined how the activity scores from our high-throughput Sort-seq experiment correlated with direct measurement of post-transcriptional repression by flow cytometry of individual peptides. We selected 16 fragments that spanned a range of activity scores, transformed our reporter strain with these individual constructs, and measured fluorescence by flow cytometry. We observed a quantitative, monotonic relationship between high-throughput activity scores and the individual fluorescence measurements, indicating that our Sort-seq approach accurately captures post-transcriptional regulation **(Figure 1F)**. This high-throughput functional profiling approach therefore provides a quantitative inventory of disordered regions that regulate gene expression post-transcriptionally and can be used to elucidate functional elements across the yeast proteome.

### Identification of post-transcriptional regulatory regions within disordered regions

We next asked how the repressive activity of disordered peptides aligned with the activity of full-length proteins. Our comparison between Sort-seq activity scores and individual fluorescence measurements suggests that activity scores below -1 correspond to fragments that decrease the YFP/RFP ratio **(Figure 1F)**. We therefore used this as a threshold to classify repressors and found 900 repressive disordered peptides derived from 395 different proteins **(Figure 2A and Figure S2A-B)**. Gene ontology analysis of proteins containing repressive fragments showed a substantial and significant enrichment in functional annotations such as deadenylation-dependent decay, mRNA destabilization, and negative regulation of translation, indicating that analysis of disordered peptides captures the regulatory effects of known full-length post-transcriptional repressors **(Figure 2B and S2C)**. Furthermore, repressive peptides were derived from high-confidence RNA binding proteins, as identified by RNA-protein interactome studies **(Figure S2B)**. Overall, this analysis indicates that our high-throughput functional profiling of disordered regions captures post-transcriptionally regulatory effects within full-length proteins.

**Figure 2:**
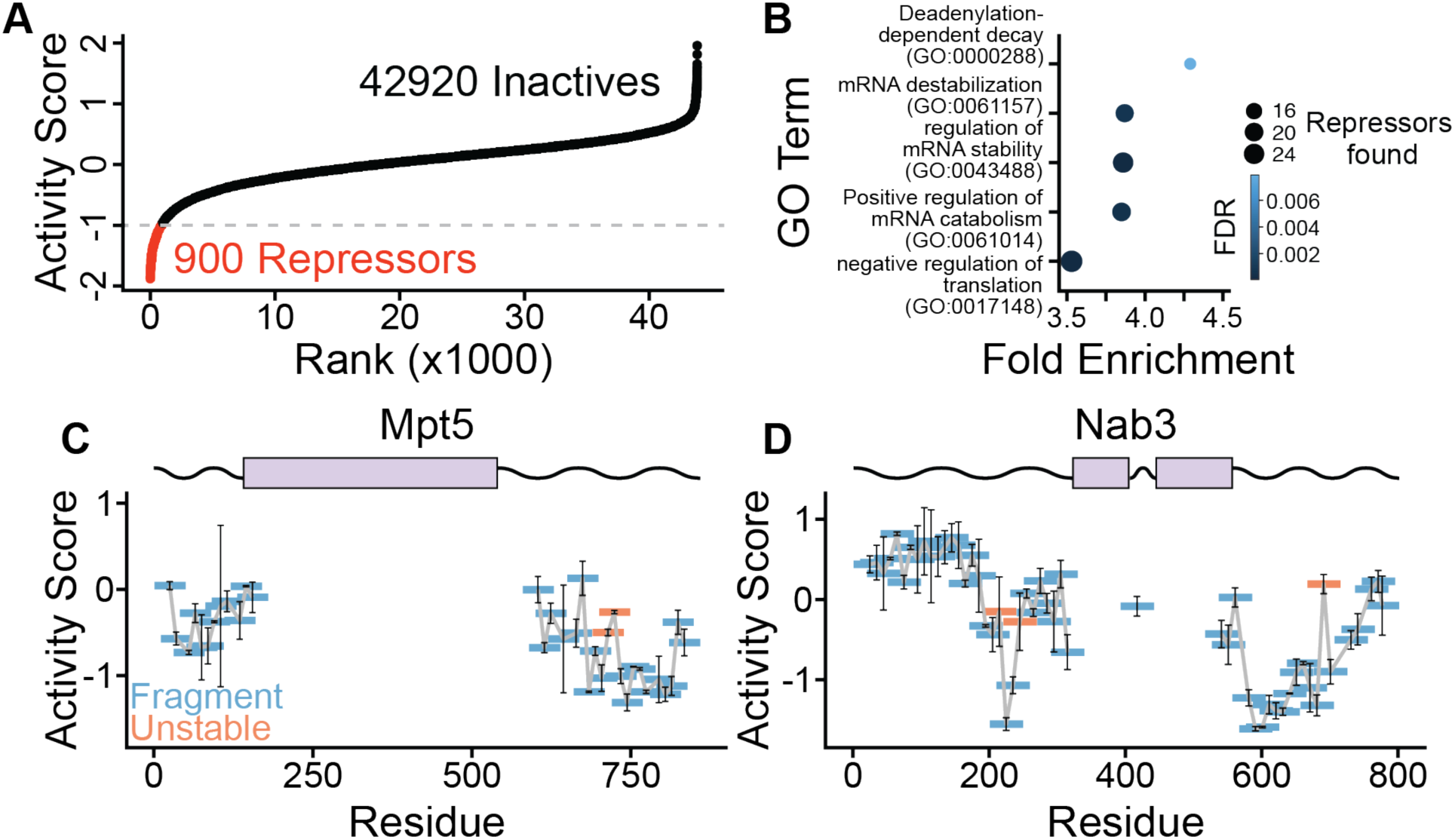
Identification of regulatory disordered regions within post-transcriptional repressors. **(A)** Rank order of average activity scores for disordered peptides (n=2). **(B)** Gene ontology enrichment analysis of proteins containing repressive fragments compared to all proteins in the disordered screening library. Select terms with fold enrichment > 3.5 are displayed with Fisher’s exact test and Benjamini-Hochberg corrected significance (*q* < 0.05). **(C)** Activity scores of fragments from Mpt5. Stable fragments are displayed in blue, unstable fragments (stability score ≥ -1) are shown in red. Standard deviation is displayed as error bars (n=2). Pfam domain boundaries are shown above **(D)** As in **(C)**, but for Nab3.

Our densely tiled peptide library further allowed us to identify functional disordered subregions within full-length regulatory proteins. We examined the fragments within each protein to find overlapping peptides with repressive activity and reconstructed functional disordered subregions. Importantly, the stability scores allow us to exclude unstable fragments that lack activity simply due to poor expression and provides higher confidence in truly repressive disordered regions **(Figure 2C-D)**. For example, we examined the yeast Pumilio and FBF family (PUF) protein, Mpt5, a known post-transcriptional regulator that binds mRNAs through its structured Pumilio homology domain and interacts with deadenylation machinery to promote turnover of target transcripts^32^. We obtained broad coverage of fragments from the disordered regions of Mpt5 and uncovered a candidate regulatory region within its C-terminus **(Figure 2C)**. These fragments fall outside the folded RNA-binding Pumilio domain and underscore how these disordered regions can recapitulate function of the full-length protein when tethered to an mRNA target. We also investigated disordered regions within Nab3, which functions in the degradation of cryptic, unspliced transcripts by the nuclear exosome^33,34^. We find two repressive subregions within Nab3 containing strong repressive activity, which may be involved in recruiting downstream mRNA decay factors to promote transcript turnover **(Figure 2D)**. Functionally profiling the disordered proteome enables us to uncover repressive subregions within longer disordered sequences and elucidate how these elements contribute to the activity of post-transcriptional regulators.

### Regulatory disordered regions function through 5′-3′ decay

Like many regulatory RNA-binding proteins, Mpt5 and Nab3 lack catalytic activity and instead recruit nucleases that degrade target mRNAs^8^. Indeed, many different post-transcriptional regulators contain short linear motifs that directly bind 5′-3′ mRNA decay machinery to promote mRNA turnover, and these nucleases are likely partners underlying the activity of disordered repressors^35–39^. We thus hypothesized that the isolated peptides in our library interact with downstream effectors to modulate YFP expression, and wanted to determine which molecular pathways these disordered regions engage to repress mRNA expression. General, or bulk 5′-3′, mRNA degradation in eukaryotes proceeds through a multistep process that begins with trimming of the poly(A) tail by the CCR4/NOT deadenylase, followed by removal of the 5’ m^7^G cap by the decapping enzyme Dcp2 and subsequent 5′-3′ exonucleolytic digestion **(Figure 3A)**^40^. We reasoned that we could learn which repressors rely on each step of this pathway by repeating our high-throughput activity profiling measurements after disrupting key components of the mRNA decay machinery and determining which peptides lost activity under these conditions. To conditionally deplete these nucleases, we individually tagged Dcp2 with the miniature auxin inducible degron (mAID) in our reporter strain and coexpressed the plant F-box protein, Tir1, which promotes ubiquitination and subsequent proteasomal degradation of mAID fusions upon treatment with indole-3-acetic acid (IAA)^41^. Likewise, we individually tagged each of Pop2 and Ccr4, the two deadenylases within the CCR4/NOT complex, with mAID. Addition of IAA drove acute depletion of the tagged protein within 4 hours and resulted in growth defects relative to the DMSO treated control, indicating functional loss of the tagged protein **(Figures 3B and S3A-E)**.

**Figure 3:**
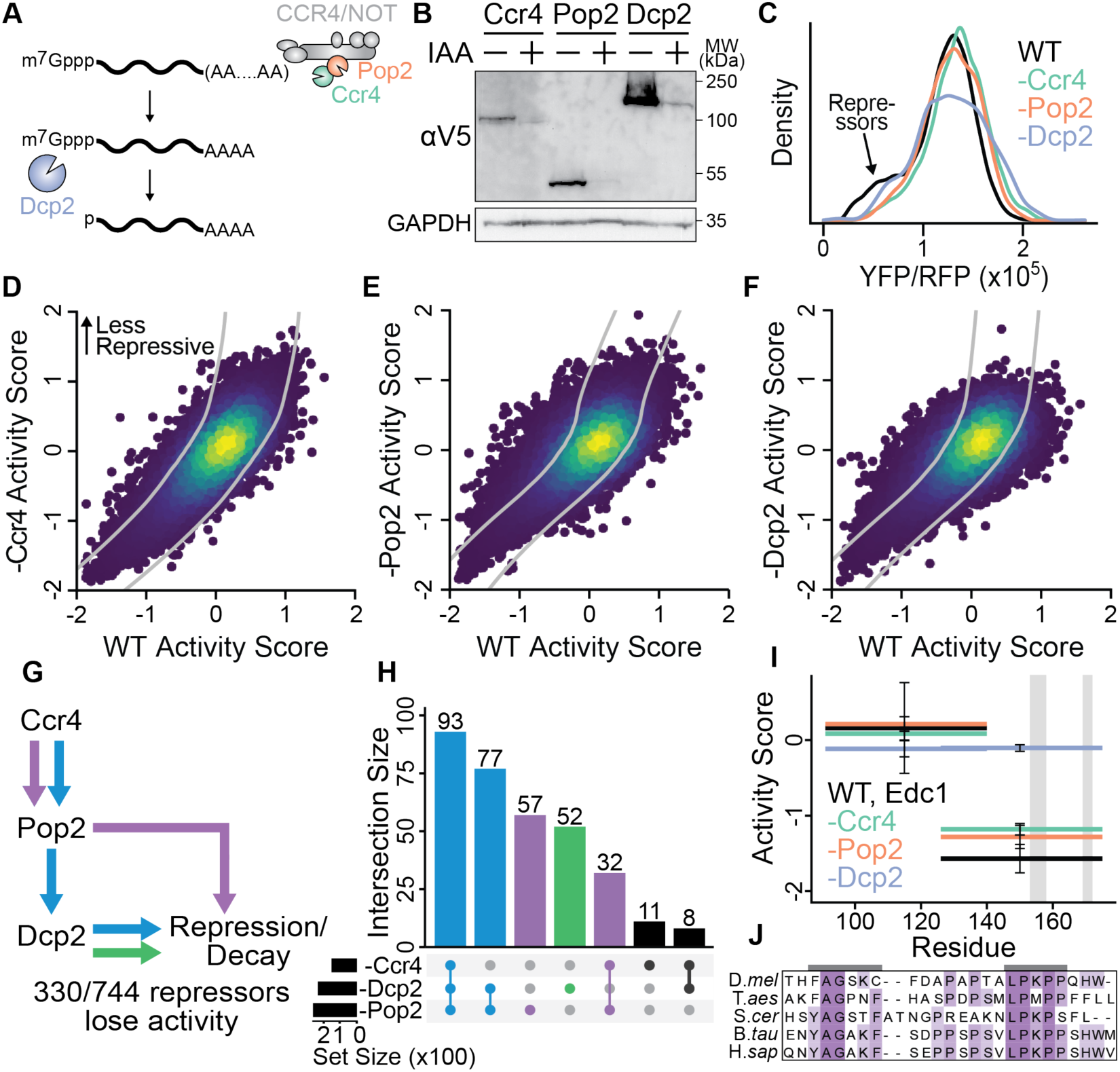
Disordered post-transcriptional regulators function through 5′-3′ mRNA decay. **(A)** Schematic of 5′-3′ mRNA decay, with deadenylation followed by decapping and exonucleolytic digestion. **(B)** Western blot of mRNA decay factor depletion and GAPDH loading control. Total protein was isolated after 4 hr of 500 µM IAA treatment or 1% DMSO control. **(C)** Fluorescence ratio distribution of degron strains expressing the full tethering library after ∼16 hour IAA treatment. The sub-population with low YFP/RFP ratio is labelled to indicate where repressors were found in the wildtype screen. **(D-F)** Comparison of average activity scores in the wildtype strain and average activity scores in strains depleted of **(D)** Ccr4, **(E)** Pop2, and **(F)** Dcp2 (n=2). Gray lines demarcate statistical significance (see materials and methods, p_adj_ ≤ 0.05). **(G)** Schematic of possible pathways leading to repression or decay of the reporter mRNA. Blue: deadenylation-dependent decay, purple: decapping-independent decay, green: direct decapping without deadenylation. Total number of repressors with significant change in activity in any degron strain shown below. **(H)** UpSet plot showing repressors that change activity following mRNA decay factor depletion. Colors as in **(G)**. **(I)** Activity scores for select fragments from Edc1 in wildtype and decay factor depletion with standard deviation displayed (n=2). Gray vertical bars highlight positions of regulatory motifs. **(J)** Sequence alignment of Edc1 motifs from yeast to metazoans.

We performed our tethering Sort-seq assay in yeast depleted of Dcp2, or either of the individual deadenylases Pop2 or Ccr4. High-throughput measurements of regulatory activity following nuclease depletion indeed revealed that many disordered peptides required 5′-3′ mRNA decay machinery for their activity **(Figure 3C-F)**. First, the bulk fluorescence ratio distribution showed a decrease in the fraction of cells with lower YFP/RFP ratio when decapping or deadenylation enzymes were depleted, suggesting a reduction in activity of some repressive fragments **(Figure 3C).** Second, comparing the activity scores in the depletion conditions to the wildtype background revealed many repressors with significantly reduced activity in each nuclease depletion condition (p_adj_ ≤ 0.05) **(Figure 3D-F)**. The activity scores between each replicate were highly correlated even in these slow growing cells **(Figure S3F-H)**. Importantly, repressors exclusively lost activity in the depletion backgrounds, while none gained activity, indicating that many of these fragments genetically depend on these nucleases for function.

We sought to classify repressive peptides based on their genetic requirements and, when possible, assign them to specific substeps of mRNA decay. We recovered 744/900 (83%) repressors with sufficient coverage in both wildtype cells and all three depletion strains. Among these peptides, 330 (44%) had significantly reduced activity in at least one depletion strain, indicating many of these repressive fragments depend on 5′-3′ decay machinery to promote mRNA turnover **(Figure 3G)**. A majority of these repressors (170/330) lost activity upon depletion of Dcp2 and at least one deadenylase, suggesting that these peptides require intact deadenylation and decapping for function **(Figure 3G-H)**. The prevalence of this pattern is consistent with idea that most mRNA decay proceeds by deadenylation followed by decapping. We also found many repressors (89) that lost activity in the Pop2 or Ccr4 depletion backgrounds but were insensitive to Dcp2 levels, indicating these fragments only require deadenylation for activity. Notably, the Ccr4 nuclease is linked to the Not1 module through Pop2; therefore, many repressors that rely directly on Ccr4 should also have reduced activity when Pop2 is depleted^42^. Supporting this, 126/145 (87%) of repressors that lost activity in the Ccr4 depletion strain also had significantly reduced function in the absence of Pop2, reflecting the structural architecture of the CCR4/NOT complex. We also observed many fragments (52) that retained activity when the deadenylases were depleted but were sensitive to Dcp2. Illustrating this dependency, we found a fragment from the Enhancer of mRNA Decapping, Edc1, which is known to directly recruit and activate the decapping complex through defined short linear interaction motifs, required only Dcp2 for its activity^43,44^ **(Figure 3I-J)**. Collectively, this analysis uncovers how numerous post-transcriptional regulatory disordered regions converge on 5′-3′ mRNA decay machinery to enforce their regulatory effects.

### Identification of repressive motifs within disordered regions through mutational scanning

Because we found that many disordered regions require 5′-3′ mRNA decay machinery for their repressive activity, we reasoned that these peptides may contain motifs that directly engage decay factors. To experimentally identify such regulatory motifs within repressive fragments, we performed mutational scanning. We selected all repressors (activity score ≤ -1) identified in wildtype cells, excluded redundant peptides that overlapped by > 30 amino acids, and tiled each fragment with a 5 amino acid GGSSG linker in one residue increments **(Figure 4A)**. We chose this longer substitution to ensure ablation of any functional elements, as some motifs may tolerate single-amino acid mutations. We transformed the reporter tethering strain with this library and performed Sort-seq as described above. We measured both activity scores, to determined how mutations affected repressor function, and stability scores, to distinguish truly inactivating mutations from those that just destabilize the tethering construct. As expected, the fluorescence ratio distribution of this library was more uniform than the wildtype library, as these fragments were derived from a highly repressive population **(Figure S4A)**. Repressor activity agreed well between the proteome-wide library and the focused mutational screen, with the 20 inactive controls among the least active, and both activity and stability scores were highly reproducible between replicates **(Figure 4B & S4B-D)**.

**Figure 4:**
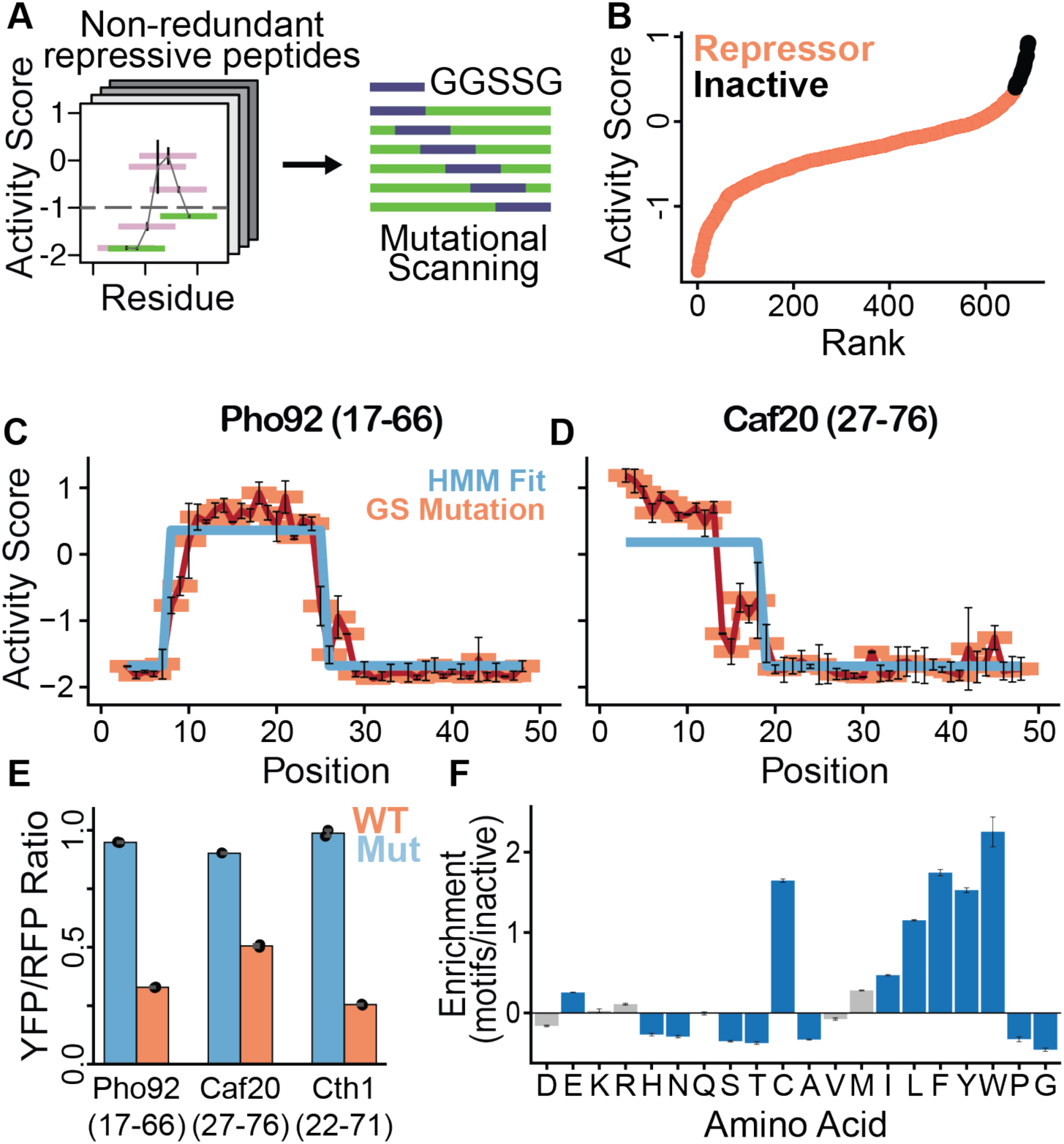
Mutational scanning of repressive disordered regions reveals functional motifs. **(A)** Schematic of mutational scanning library design. Each repressor was tiled with a GGSSG linker in one amino acid increments. For repressors that overlapped by > 30 amino acids, only the most active fragment was included in the mutational library (679 repressors). **(B)** Rank order of average activity scores for the wildtype peptides in the mutational library, with repressive fragments shown in red and 20 inactive controls in black (n=2). **(C)** Mutational scanning profile of repressive Pho92(17-66) peptide. Average activity scores for each mutational window are shown in light red, with standard deviation, connected by dark red line (n=2). Hidden Markov Model fits for the mutational scans shown in blue. Position indicates mutational window within the fragment. **(D)** As in **(C)**, but for Caf20(27-76). **(E)** Flow cytometry of wildtype repressors and mutations expected to disrupt activity in red and blue, respectively. Mutations are Pho92(17-66)-Mutation window 16, Caf20(27-76)-Mutation window 2, and Cth1(22-71)-Mutation window 22. Mean YFP/RFP ratios of individual replicates and standard deviation are displayed (n=2). **(F)** Enrichment of amino acids within mutationally sensitive regions compared to regions that tolerate the GGSSG substitution, with standard deviation shown for 10,000 bootstrap iterations and colored blue if significant (uncorrected p-value < 0.05).

We next wanted to use the mutational scanning scores to identify repressive motifs within disordered regions. We reasoned that we could classify motifs by finding a series of contiguous mutations that all abolished repressor activity. To objectively quantify these regions and discover motifs, we fit the mutational scanning profiles of each repressor to a two-state Hidden Markov Model (HMM). We found that 230/665 repressors had statistical support for an HMM with two different activity levels across mutant peptides (p_adj_ ≤ 0.05) and generated maximum likelihood HMM profiles to visualize mutationally sensitive regions within each repressor. For example, the HMM clearly identified a mutationally sensitive region with a strongly repressive peptide from the yeast YTH domain containing protein, Pho92 (residues 17-66), that may be a repressive motif **(Figure 4C)**^45^. Additionally, we found a mutationally sensitive region within a fragment from one of the yeast eIF4E binding proteins, Caf20 **(Figure 4D)**. These residues map to a helix of Caf20 that directly contacts eIF4E in the experimentally determined crystal structure of eIF4E-Caf20 complex, highlighting the ability of this mutational scanning approach to uncover functionally important residues within repressive disordered regions^46^. We selected three repressive fragments, introduced mutations expected to disrupt their activity based on the HMM analysis, and measure their activity by flow cytometry. In all cases, these mutations abrogated repressive activity of the parent peptide **(Figure 4E)**. Thus, this mutational tiling approach enables identification of functionally important residues within repressive disordered regions.

This HMM analysis allows us to define motifs at single-amino acid resolution across our diverse library of repressors. We wanted to learn the sequence features of these motifs that distinguish them from inactive disordered regions. We therefore compared the amino acid composition of residues found within the motifs to those outside the mutationally sensitive regions. Motifs were enriched in aromatic and large aliphatic residues relative to the background, suggesting that such residues may be functionally required for repressor activity **(Figure 4F)**. Furthermore, this pattern is consistent with these motifs forming protein-protein interactions with mRNA decay factors through buried hydrophobic residues^6^. Interestingly, prolines were depleted in the motifs relative to the flanking, inactive residues, which may indicate that these regions fold upon interacting with their binding partners. Systematically defining functional elements within disordered regions allows us to learn the molecular rules governing how repressive post-transcriptional regulatory motifs promote mRNA turnover and translation repression.

### Repressive motifs drive activity of post-transcriptional regulators

We next wanted to understand how these motifs discovered by peptide mutational scanning contributed to regulatory activity in the context of their full-length protein. We first focused on the tandem CCCH-type zinc finger protein Tis11, which promotes degradation of mRNAs containing AU rich elements (AREs) in response to Fe(II) depletion^47^. Our mutational scanning data identified a functional motif within Tis11 and captured substitutions that abolished function **(Figure 5A-B)**. Additionally, sequence alignment between *S. cerevisiae* Tis11 and homologues from other yeast species found a conserved motif within the N-terminal disordered region **(Figure 5C)**^48^. The human ortholog of Tis11, TTP/ZFP36, is known to interact with CNOT1 (Cdc39 in yeast), and computational modelling of this Tis11 motif with Cdc39 revealed a conserved interaction **(Figure 5D)**^37^. To assess how this motif contributes to the function of Tis11, we selected a mutation that crippled activity of the isolated Tis11 fragment in our tethering assay, introduced this into the endogenous *tis11* locus, and measured how mutant yeast responded to iron depletion. Disruption of the motif in *tis11* phenocopied the severe growth defect of the whole-gene deletion when iron was depleted from the medium with ferrozine, suggesting that the motif is required for Tis11-mediated post-transcriptional regulation **(Figure 5E)**. This effect was not due to decreased protein abundance and could be fully reversed upon reintroduction of Fe(II) to the media **(Figure 5SA-C)**, demonstrating that mutational scanning of regulatory peptides can uncover essential motifs required for the activity of endogenous post-transcriptional regulators.

**Figure 5:**
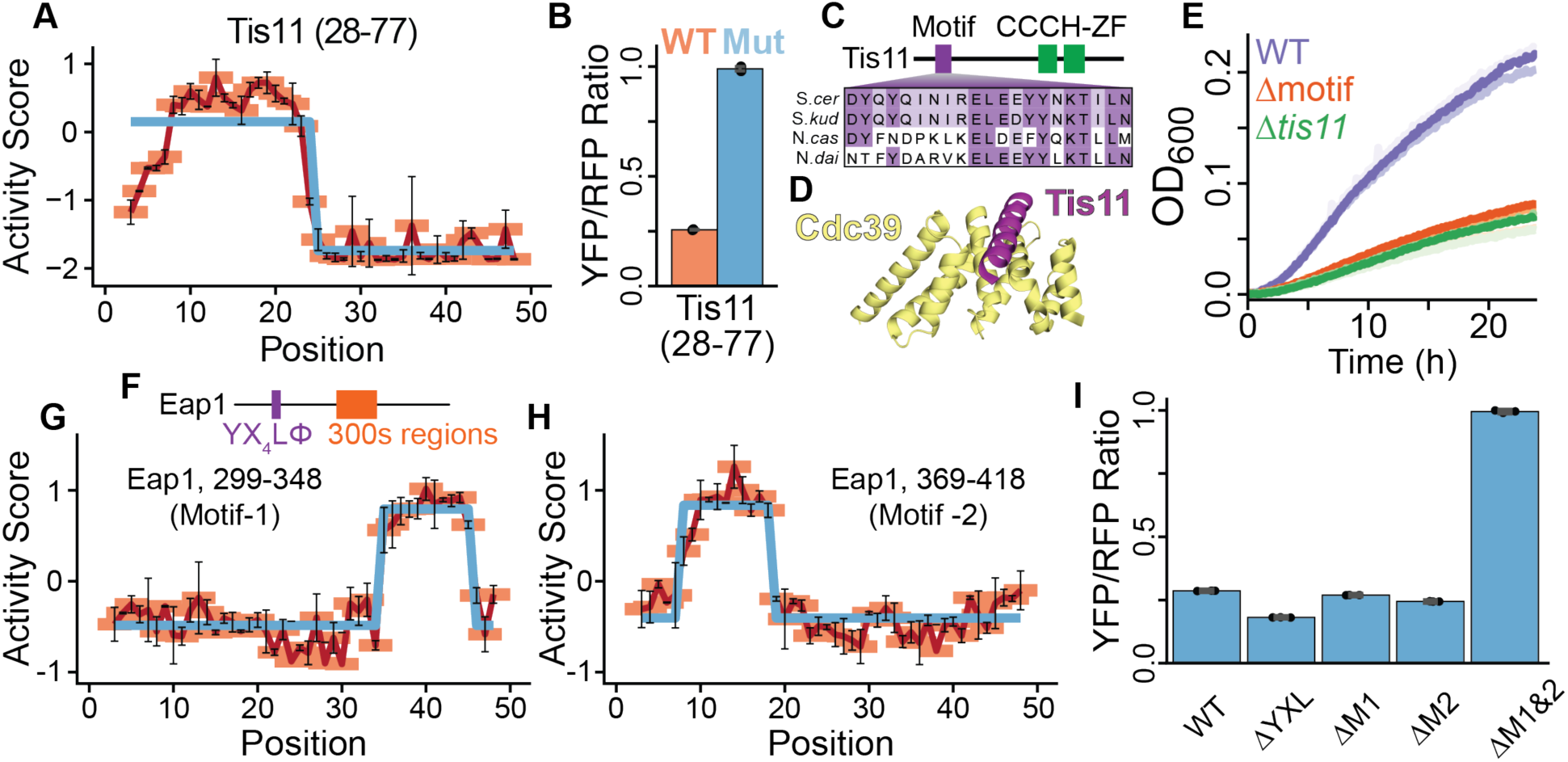
Repressive motifs identified by mutational scanning are required for post-transcriptional repressor function. **(A)** Mutational scanning profile of Tis11(28-77), as in Figure 4C. **(B)** YFP/RFP ratio of Tis11(28-77) and Tis11(28-77)-Mutation11 in red and blue, respectively. YFP/RFP ratios of individual replicates and standard deviation are displayed (n=2). **(C)** Schematic of Tis11 showing the CCCH-Zinc Finger (green) and motif (purple). Sequence alignments for representative yeast displayed below. **(D)** AlphaFold2 model of Tis11 motif and the mIF4G domain of Cdc39/Not1. **(E)** Growth curves of yeast expressing different V5-tagged Tis11 constructs, and Δ*tis11* (n=3) in complete synthetic media with 750 µM ferrozine. Three biological replicates are displayed in different shades. **(F)** Gene diagram of Eap1, with the Y(X_4_)Lφ and repressive motif regions shown in purple and red, respectively. **(G-H)** Mutational scanning profiles for motifs within Eap1. Residues indicated in the panel legend. **(I)** YFP/RFP ratio of reporter yeast expressing different full-length Eap1 constructs containing mutations as indicated. ΔYXL replaces the Y(X_4_)Lφ motif (residues 126-132) with GGSSG, ΔM1 replaces residues 332-342 and ΔM2 replaces residues 376-386 with GSG linkers, and ΔM1&2 combines ΔM1 and ΔM2. Mean fluorescence ratios of biological replicates are shown along with standard deviation (n=3).

As another example, our mutational scanning data identified two conserved fungal motifs within a second eIF4E-binding protein, Eap1. Like Caf20, Eap1 is known to bind eIF4E using the conserved Y(X_4_)Lφ motif on the same surface as eIF4G and thus inhibits formation of the eIF4F complex required for translation initiation **(Figure 5F)**^46,49,50^. The Y(X_4_)Lφ motif was not included in our screen and is dispensable for Eap1 function *in vivo*^51,52^. However, our mutational scanning approach revealed two repressive motifs within Eap1 in residues 331-341 and 377-384. These motifs were distal from the Y(X_4_)Lφ motif and conserved in related yeast species **(Figure 5G-H and S5D-E)**. Tethering full-length Eap1 reduced reporter expression, and deletion of the Y(X_4_)Lφ motif did not affect repression by tethered Eap1. In contrast, deleting the motifs we originally uncovered by peptide mutational scanning abolished the ability of full-length Eap1 to repress reporter expression. Furthermore, these motifs functioned redundantly, as Eap1 constructs with individual motif mutations still decreased YFP output **(Figure 5I)**. These newly identified motifs appear to act independently from the Y(X_4_)Lφ motif to repress translation, consistent with the dispensability of the Y(X_4_)Lφ motif^51,52^. By cataloging motifs through mutational scanning, we can reveal how these elements underlie the regulatory activity of full-length post-transcriptional repressors.

### Overall sequence composition drives repressive activity of disordered regions

In addition to repressive disordered regions harboring mutationally sensitive motifs, we also observed many peptides that retained activity across all tested mutations. For example, no mutations in the peptide derived from residues 151-200 of the translation repressor Sgn1 significantly disrupted activity **(Figure 6A)**. This mutational robustness could be explained by an individual motif that is not sufficiently crippled by a 5 amino acid substitution, or multiple, redundant motifs dispersed throughout the repressor. Alternately, these peptides may depend only on their overall physicochemical properties for repressive function. To distinguish these possibilities, our mutational scanning library also included five random permutations of each repressive peptide sequence. Scrambling should disrupt long or redundant motifs, but will not alter the overall sequence composition of the peptide. We computed an average activity score across the scrambled variants for each repressor and found that 65% (435/665) of fragments had similar activity between the wildtype and shuffled sequences **(Figure 6B)**. This tolerance to permutation indicates that these fragments may rely on overall sequence composition to repress reporter mRNA expression.

**Figure 6:**
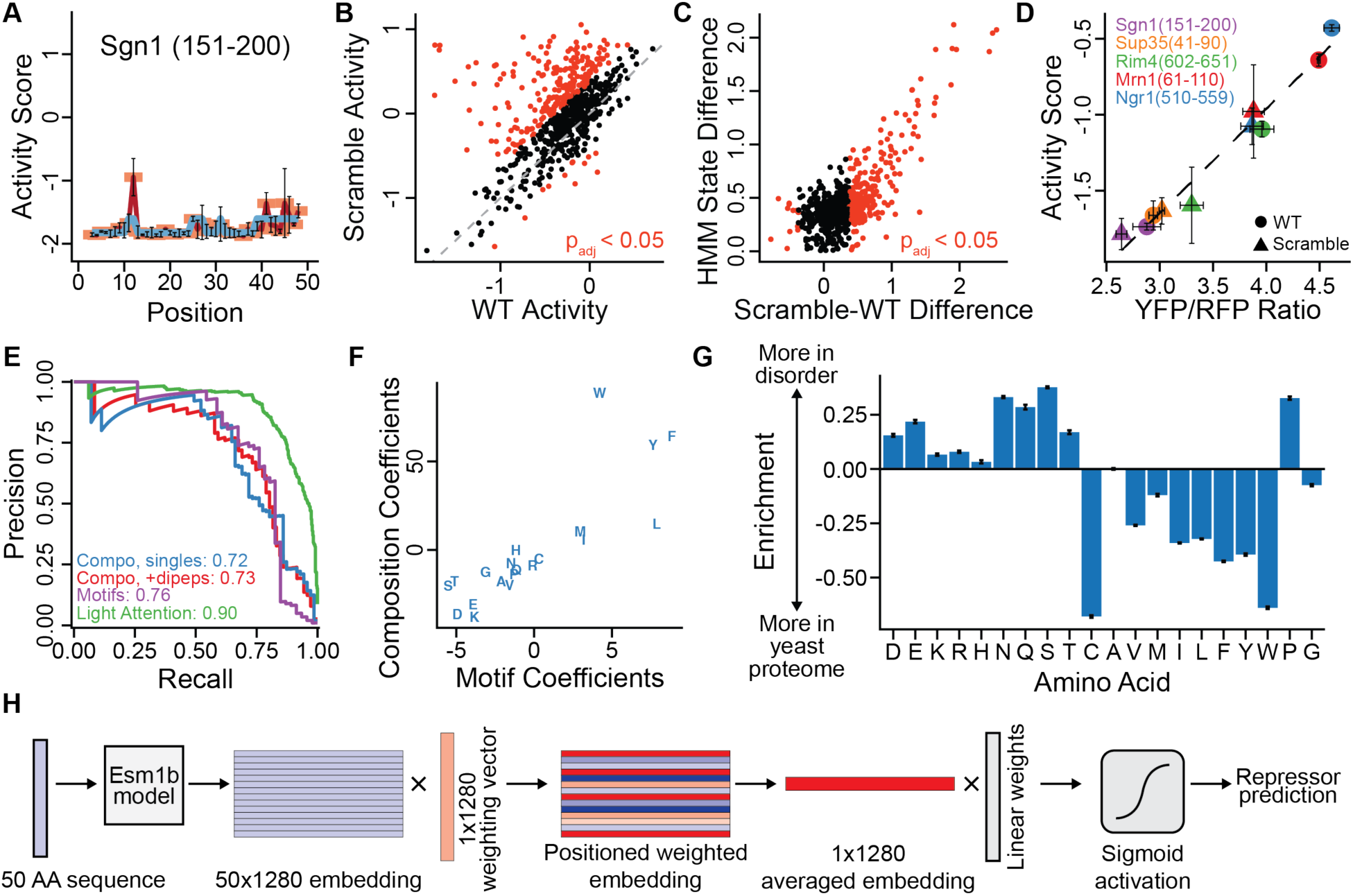
Aromatic amino acids are enriched in repressive disordered regions. **(A)** Mutational scanning profile of Sgn1(151-200), as in Figure 4C. **(B)** Activity scores for each wildtype peptide plotted against the average of the associated scrambles. Wildtype activity scores are derived from the HMM fits to the wildtype repressor. Red points are significantly different (Benjamini-Hochberg corrected p-value ≤ 0.05). **(C)** Correlation between HMM state differences in mutational scanning and scramble differences in activity score. Points colored as in **(B)**. **(D)** Correlation between average activity scores from mutational scanning (n=2) and individual YFP/RFP ratios by flow cytometry with standard deviation (n=3 for flow cytometry, n=2 for activity scores). Dashed line shows fit of linear model. **(E)** Precision-recall curves for models predicting repressor activity from peptide sequence. The light attention model was trained only on composition-dependent fragments. AUPRC displayed in panel legend. **(F)** Coefficients of the L2-penalized logistic regression models for composition-driven and motif-dependent models using primary amino acid sequence. **(G)** Enrichment of amino acids within the disordered screening library compared to the yeast proteome, with standard deviation of 10,0000 bootstrap iterations. **(G)** Schematic showing how protein language representations are coupled with a light attention classifier to predict functional residues within repressors.

Repressors that retain activity when shuffled are fundamentally distinct from motif-containing fragments, which require the correct linear sequence of amino acids to engage binding partners. One prediction of this difference is that mutational substitution should not affect the activity of composition-driven, scramble-insensitive repressors, whereas peptides with mutationally sensitive motifs should be inactivated by scrambling. Indeed, repressors with a larger activity difference between functional and inactive mutational variants were universally more sensitive to sequence permutations **(Figure 6C)**. This correlation confirms that peptides harboring defined motifs lose activity when these residues are randomly re-distributed across the peptide sequence. Thus, we used this scramble sensitivity to separate repressors into two distinct classes: motif-dependent fragments that lose activity when shuffled, and composition-driven repressors that retain activity even when permutated. We then selected 5 composition-driven repressors, along with an associated scramble for each, and measured their effects on reporter expression by flow cytometry. In all cases, the scrambles were at least as active as their wildtype counterpart, with excellent agreement between the activity scores from the high-throughput approach and the individual measurements **(Figure 6D)**. This robustness stands in contrast to the mutational sensitivity of motif-containing fragments **(Figure 4E & 5B)**. These permutations imply regulators use two distinct classes of functional disordered regions with different linear sequence requirements to repress gene expression post-transcriptionally.

### Aromatic amino acids are associated with regulatory activity in disordered regions

After identifying two types of repressors that depended on either motifs or overall amino acid composition, we wanted to learn what sequence features were required for activity within each class. Taking advantage of the scale of our high-throughput data, we constructed logistic regression classifiers that predicted repressors using only the primary amino acid sequence, as these provided permutation-independent features for composition-driven repressors. We used the scramble sensitivity to divide repressors into either composition-driven or motif-dependent fragments and separately modelled the amino acid composition of isolated repressive motifs and composition-driven repressors, contrasted with inactive disordered peptides. In all cases, these models achieved high predictive accuracy (AUPRC 0.72-0.76), suggesting the primary amino acid sequence was indeed a major determinant of regulatory activity in disordered regions **(Figure 6E)**.

We then inferred which sequence features were associated with repressor activity by examining the weights of our logistic regression models. Aromatic amino acids were strongly associated with both motif-dependent and composition-driven repressors, while acidic residues and lysine were negatively correlated with repressive activity **(Figure 6F)**. Supporting these relationships, peptide activity increased with the number of aromatic amino acids and decreased with number of negatively charged residues within each sequence **(Figure S6A-B)**. Aliphatic residues were also associated with post-transcriptional repressive activity, though to a lesser extent than the aromatic amino acids **(Figure 6F)**. Incorporating dipeptide features only marginally improved predictive power, suggesting that residues driving repressor activity do not strongly depend on their immediate neighbors **(Figure S6C)**. While hydrophobic and aromatic residues were associated with activity, there was no correlation between overall hydrophobicity and repressor activity; this uncoupling indicates that additional properties contribute to function **(Figure S6D)**. However, the activity of composition-dependent repressors was correlated with the degree of dispersion of aromatic amino acids throughout the sequence and may imply that some patterning of these residues is required for activity **(Figure S6E)**^53^. Our interpretable logistic models thus provide insights into the molecular grammar of post-transcriptional regulatory disordered regions.

We next tested whether our logistic models could distinguish between these two classes of repressive disordered regions. The surprisingly strong correlation between the logistic regression coefficients for motif-dependent and composition-driven repressors indicate that similar amino acids are associated with both classes of regulatory disordered regions **(Figure 6F)**. This relationship implies that, despite different mutational and scrambling tolerances, both classes of repressors rely on similar amino acid features. However, these models could not strongly predict the other class, suggesting that additional properties from other amino acids may contribute to each type of repressor **(Figure S6F)**. The importance of aromatic residues for repressor function was striking, given that these residues are typically depleted from disordered regions. Accordingly, aromatic amino acids and hydrophobic residues were underrepresented in our disordered library relative to the yeast proteome **(Figure 6G).** These models imply that aromatic-rich elements within disordered regions may influence the post-transcriptional regulatory activity of repressors, regardless of the presence of conserved motifs.

### Predicting repressive disordered regions using protein language models

While overall amino acid content predicted repressor function well, it necessarily treats each occurrence of an amino acid equally and cannot identify functionally important positions within a sequence. Furthermore, the remaining gap in predictive performance suggests that additional features, such as the context or patterning of amino acids within a peptide, are also important for repressor activity. We thus sought to predict regulatory peptides using protein language models, which capture contextual information about each residue of a protein sequence in latent space representations^54^. While these representations are not directly interpretable, they support remarkably good predictions of protein structure and function with no additional tuning of the protein language model itself (i.e., zero-shot learning)^55^. We reasoned these representations could help identify functionally important positions within composition-driven repressors. Towards this end, we aimed to a build a repressive peptide predictor that used these protein language models to categorize composition-driven repressors and identify functionally important residues within them.

To accomplish this task, we trained a light attention model to distinguish composition-dependent repressors from inactive disordered peptides **(Figure 6H)**^56^. Starting from ESM-1b latent space representations of peptides that treat each residue uniquely depending on its context within the fragment, our model learns both a weighting function that emphasizes residues associated with activity and a scoring function for the weighted average that predicts overall repressive peptides. Crucially, the weighting function provides a positionally equivariant weight that recognizes residues that contribute to repression wherever they occur within a sequence, in the context of a model that scores the overall activity of the fragment. Our light attention model substantially improved predictive power (AUPRC = 0.9) relative to classifiers using amino acid content alone, demonstrating the value of coupling these protein language representations to predictive models **(Figure 6E)**. In addition to increasing performance, this light attention architecture allows us to directly identify important residues within a repressor by examining the weighting layer. As shown in two representative cases, this model often highlighted aromatic amino acids—the same residues predicted to be important by the logistic regression approaches **(Figure S6G-H)**. However, not every aromatic residue carried equal weight, suggesting that local physicochemical properties or context may affect how a specific residue contributes to repressor activity. Furthermore, while logistic models treat scrambled sequences equally, this position-aware model can find residues within permutations that are associated with function, consistent with the positional invariance of this model **(Figure S6H-I)**. Combining protein language models with interpretable machine learning approaches allows us to understand the molecular grammar underlying composition-driven repressors and may enable more broad exploration of sequence features used by disordered regions in diverse biological processes.

## Discussion

While the importance of disordered regions for regulating mRNA stability and translation is well appreciated, the molecular features and the pathways used by these sequences remains poorly understood. Here, we systematically discovered disordered regions that control gene expression post-transcriptionally using a high-throughput functional profiling approach. Quantitative activity measurements from massively parallel tethering libraries allowed us to evaluate the regulatory potential of the entire disordered proteome and identify functional elements within longer disordered regions. We further delineated how these peptides repress mRNA expression by testing how their activity changes when key 5′-3′ decay nucleases are disrupted. Finally, analysis of systematically mutated variant libraries enabled us to define distinct classes of disordered regions that either contain short linear motifs or require only overall sequence composition for their activity. By applying machine learning to this high-throughput dataset, we revealed how aromatic amino acids within disordered regions promote repressor activity, modified by their context and patterning. This comprehensive analysis defines the molecular principles and biochemical mechanisms used by disordered regions to regulate gene expression by modulating translation and mRNA decay.

Elucidating how regulatory proteins bind mRNA targets and influence their expression is a major challenge towards understanding post-transcriptional regulation. While it is known that many of these proteins use combinations of globular domains and disordered regions to control mRNA stability and protein synthesis, the contribution of these disordered regions to molecular function has remained unclear. By coupling regulatory activity to a fluorescence phenotype, we were able to systematically identify repressive disordered regions on a proteome-wide scale and understand which elements engender biological regulation. Hundreds of individual 50-amino acid peptides each sufficed to repress reporter expression, showing that many disordered regions require only short regulatory elements within them to enact post-transcriptional control. Furthermore, these regions retained activity independently of any globular domains when tethered to a target mRNA. The separability of disordered regions from their endogenous, structured domains in post-transcriptional regulators is analogous to the modular architecture of transcription factors, which use folded DNA binding domains to bind specific targets in the genome and then recruit transcriptional machinery through unstructured activation domains. Indeed, several studies have catalogued and dissected transcription factor activation domains that promote transcription when tethered to DNA^3–5,57–59^. Our work extends this paradigm to post-transcriptional regulators and suggests disordered regions within RNA binding proteins may likewise drive their regulatory function.

The repressive disordered regions that we identified do not possess catalytic activity on their own; instead, these fit a recurring pattern where post-transcriptional repressors recruit downstream decay factors to promote mRNA turnover^8^. Accordingly, we found that many repressive disordered peptides lost activity when key mRNA decay nucleases were depleted. The genetic dependencies of these peptides varied, indicating that different regulatory disordered regions require distinct nucleases to repress expression. This analysis allowed us to reconstruct the mRNA decay pathways used by these disordered regions and highlights how many mRNA regulators converge on a few downstream nucleases to promote mRNA turnover.

Interestingly, we could not detect any consensus motifs shared between regulatory peptides, suggesting that diverse sequences within different disordered regions confer the same function. By experimentally surveying each repressive peptide through unbiased mutational scanning, we precisely delineated dozens of unrelated motifs within repressors using quantitative HMMs. In two cases, we confirmed that the short linear motifs we identified in tethered peptides were required for the function of the full-length, endogenous protein. Repressive motifs were enriched in aromatic and hydrophobic amino acids, pointing towards a role in forming interactions with downstream effector machinery. Our comprehensive survey lays the foundation for future studies to uncover the binding partners and protein interaction networks used by these disordered regions to regulate mRNA stability and protein synthesis.

Strikingly, most repressive peptides lacked motifs and retained activity even when their sequences were scrambled. This mutational tolerance is fundamentally distinct from the sensitivity of motif-dependent repressors and suggests that these peptides instead function through overall sequence composition. Such composition-dependent repressors may engage their interaction partners through fuzzy binding modes, similar to transcriptional activation domains that recruit the Mediator complex^60^. Fuzzy interactions differ from the coupled folding often observed when linear motifs bind their partners^61^. Taken together, this analysis reveals that post-transcriptional repressors employ diverse interaction mechanisms to mediate their regulatory effects. Additionally, while motifs can often be gleaned through overall sequence conservation, composition-driven functional peptides typically lack an obvious evolutionary signature^17,62^. Our mutational scanning approach can thus identify repressors that cannot be found through conservation alone and illustrates the broad range of sequence requirements governing disordered regions that control post-transcriptional gene expression.

While we defined two classes of disordered repressors with different sequence constraints, they both share an enrichment of aromatic amino acids, which are otherwise depleted from the disordered proteome. Biased amino acid composition can thus highlight sections within longer disordered regions that may contain post-transcriptional regulatory activity. While aromatic residues were associated with repressive activity, negatively charged amino acids seemed to reduce function. The depletion of negative charges within repressors stands in contrast to the fuzzy-binding disordered regions observed in transcriptional activation domains, which require acidic residues; this distinction suggests a complex molecular grammar that can encode either transcriptional or post-transcriptional regulation—and perhaps other functions—likely through favoring interactions with specific cofactors^3,4^. Interestingly, serine and threonine were depleted from repressors; these residues can become negatively charged through phosphorylation and may be phosphoswitches that dynamically tune how these repressors interact with their binding partners.

Though motif-dependent peptides strictly require the correct linear sequence of amino acids, even composition-driven repressors are influenced by the patterning of residues within the peptide. While these sequences tolerate most mutations well, select alterations can cause striking changes in downstream activity^63^. To distinguish key functional residues, we created a light attention model that learns these additional repressive features from large protein language models. This model improved our activity predictions and provides a valuable framework for further understanding the molecular grammar of disordered sequences in broad regulatory programs, such as transcription activation domains and E3 degron signals^3,22,57,64^. These approaches also exemplify a strategy to understand the effects of sequence variants and mutations in disordered peptides, which have distinct behavior from folded domains and are less suited to structure-guided approaches. Coupling attention models with protein language representations and high-throughput analysis may help distinguish mutations, or post-translational modifications, that modulate regulatory activity of disordered regions.

In this work we have identified disordered elements that mediate post-transcriptional control of gene expression, uncovered the biochemical pathways required for their activity, and defined the molecular grammar determining their function. While we pinpoint regulatory elements within longer disordered regions, the surrounding regions likely play additional roles in modulating repressive activity, perhaps by altering higher-order interactions or phase separation behavior. Our work also raises important questions about how disordered regions and globular domains cooperate within regulatory proteins and interface with additional *trans* acting factors. Defining these connections promises to clarify the regulatory mechanisms underlying post-transcriptional repressor function. Relatedly, determining how these regulators influence endogenous mRNAs may inform on how specific transcript properties—such as RBP binding sites, translation efficiency, codon content, or initiation rates—modulate the function of post-transcriptional regulatory proteins. Understanding how these disordered elements work within broader regulatory networks will provide a complete framework for how proteins post-transcriptionally shape gene expression programs.

## Acknowledgements

We thank Hector Nolla for help with the flow cytometry and Twist Biosciences for assistance with the oligonucleotide library design. We thank Andreas Martin and Andrew S. Lyon for comments on the manuscript, Max V. Staller for helpful discussion, Izaiah Ornelas for assistance with cloning, and members of the Ingolia and Liana Lareau laboratories for suggestions throughout the study. This work was supported by NIH Ruth L. Kirschstein Postdoctoral Fellowship 5F32GM148044-02 (J.H.L) and R01 GM130996 (N.T.I). Sequencing was supported by an NIH S10 OD018174 instrumentation grant to the QB3 Genomics facility at UC Berkeley (RRID:SCR_022170).

## Author contributions

J.H.L and N.T.I designed the study. J.H.L performed all experiments and analysis, with help from N.T.I for genomics and computational approaches. J.H.L wrote the manuscript with input and editing from N.T.I, who supervised the project.

## Lead contact and materials availability

Requests for reagents and resources should be directed to and will be fulfilled by the lead contact, Nicholas T. Ingolia

---

**Figure S1:**
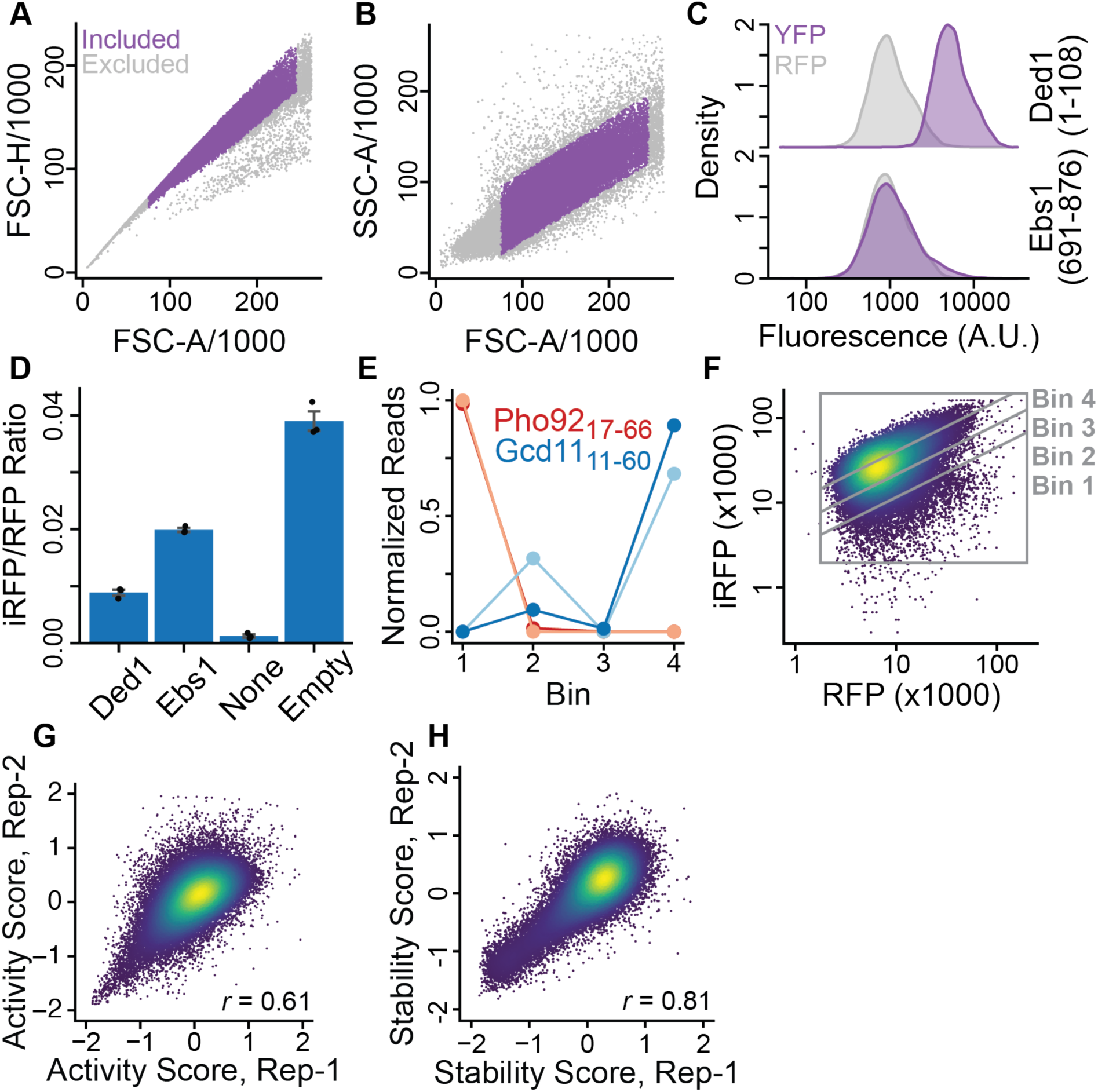
Validation of high-throughput tethering approach. **(A-B)** Gating schemes used for flow cytometry analysis. These criteria were applied across all subsequent analyses. **(C)** Representative raw YFP and RFP distributions for tethering controls. **(D)** mean iRFP/RFP ratios from flow cytometry for tethering fragments. Individual values from each replicate and standard deviation are displayed (n=3). **(E)** Normalized read distribution of two fragments across bins. Color shades show two independent biological replicates. **(F)** Gating scheme for FACS on the iRFP/RFP distribution. Bins divide the cells into four roughly equal-sized populations. **(G-H)** Correlation between replicates for **(G)** activity scores and **(H)** stability scores, with Pearson correlation coefficient displayed in figure panels.

**Figure S2:**
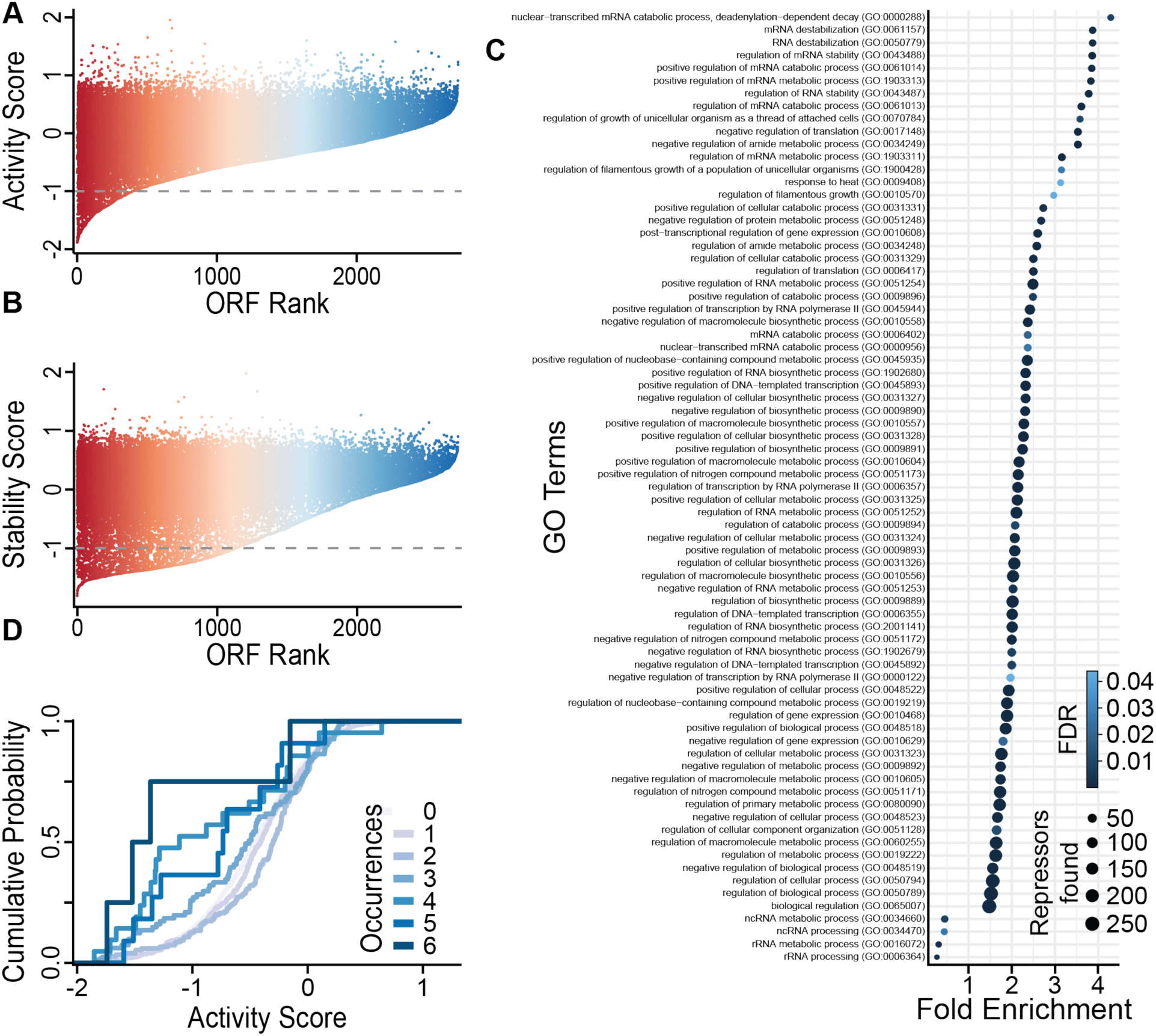
Additional analysis of disordered fragments within full-length proteins. **(A-B)** Waterfall plot of **(A)** activity scores and **(B)** stability scores for fragments within each ORF. Threshold scores used to identify repressors or unstable fragments are shown in gray. **(C)** Full gene ontology analysis of statistically significant terms for genes with repressive fragments with the Benjamini-Hochberg corrected p-values from Fisher’s exact test. **(D)** Cumulative distribution function of genes with repressive peptides found across different RNA-protein interaction datasets from yeast^9^.

**Figure S3:**
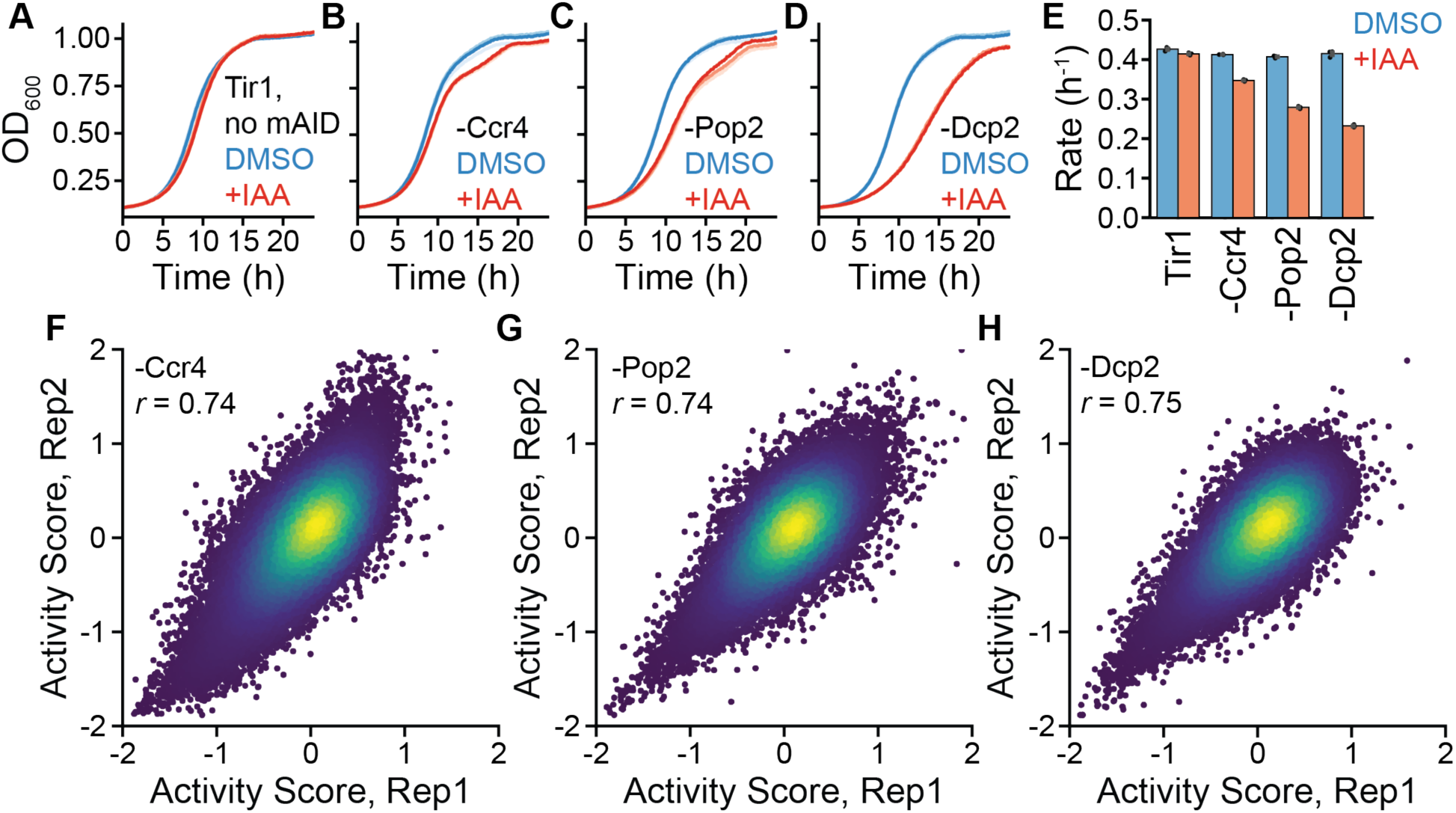
Depletion of core mRNA decay nucleases and activity scores in these backgrounds. **(A-D)** Growth curves of reporter strains expressing **(A)** the Tir1 F-box protein only, or with the mAID tag on **(B)** Ccr4, **(C)** Pop2, or **(D)** Dcp2. All strains were treated with 500 µM IAA or 1% DMSO as indicated. Different shades correspond to individual biological replicates (n=3). **(E)** Fitted growth rates for the curves shown in **(A-D)**. Mean of fits for individual biological replicates and standard deviation are displayed (n=3). **(F-H)** Correlation between activity scores for biological replicates of tethering library in depletion strains after 16 hr of IAA treatment for **(F)** Ccr4 depletion, **(G)** Pop2 depletion, and **(H)** Dcp2 depletion, with Pearson correlation coefficients shown in panel legends.

**Figure S4:**
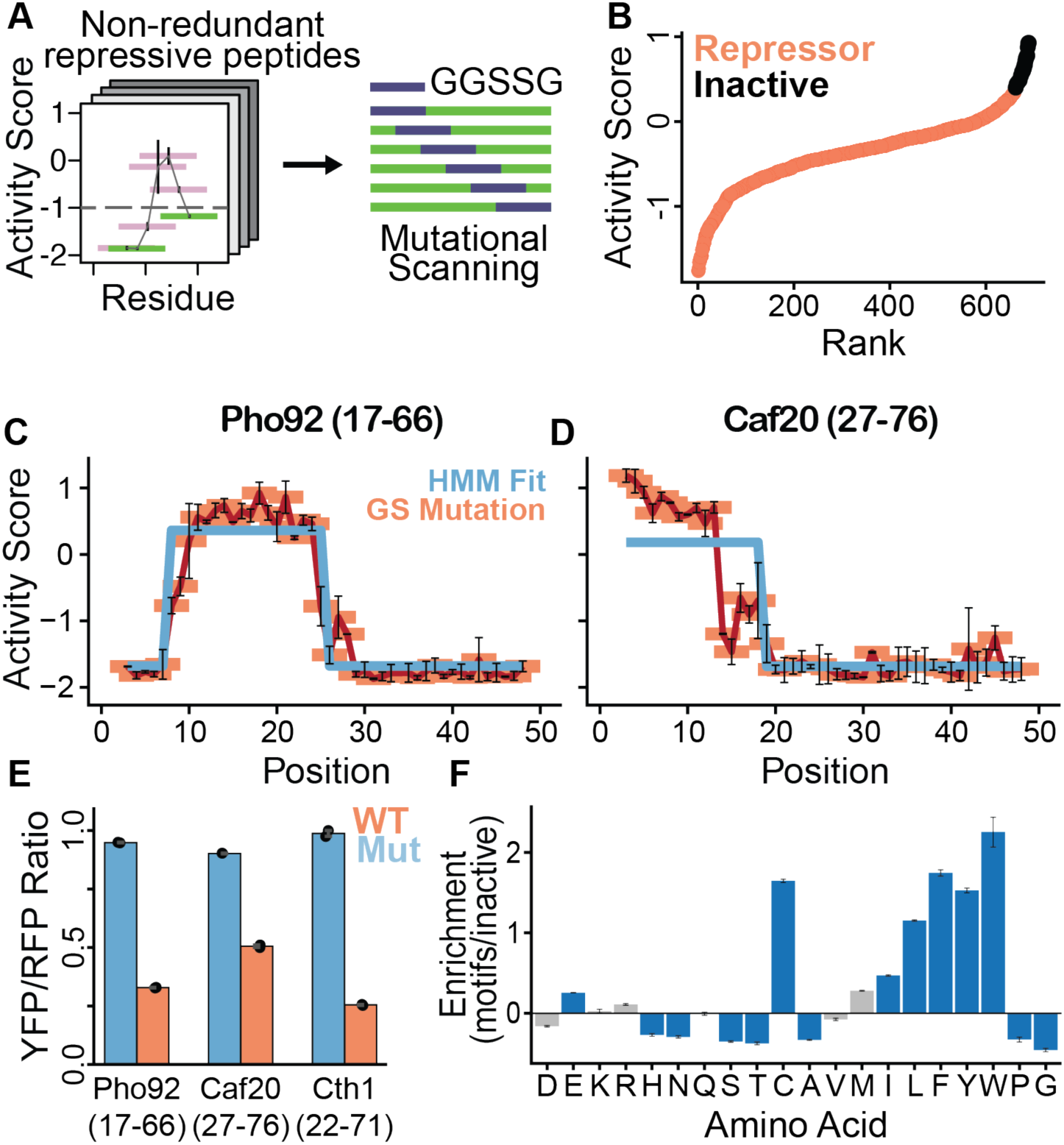
Sort-seq of mutational scanning library and activity scores. **(A)** Distribution of YFP/RFP ratios of reporter yeast expressing the mutational scanning library. Vertical lines demarcate equal population bins used in sorting. **(B-C)** Correlation between **(B)** stability scores and **(C)** activity scores for biological replicates of the mutational scanning screen. The Pearson correlation coefficient is displayed in panel legend. **(D)** Correlation between average activity scores for wildtype repressive fragments in the mutational scanning library screen and wildtype disordered proteome library screen (n=2). Repressors are shown in black and 20 inactive controls shown in red.

**Figure S5:**
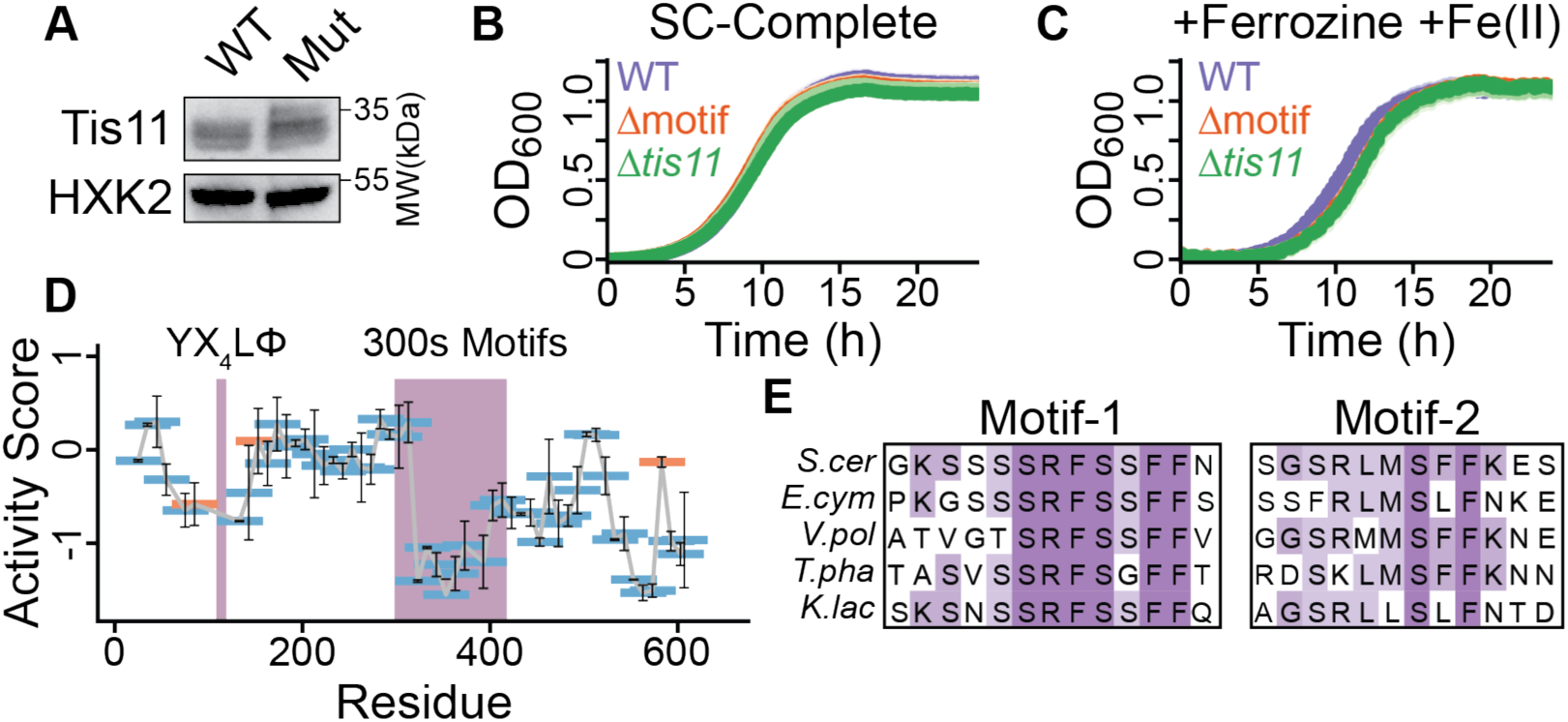
Functional analysis of motifs within post-transcriptional repressors. **(A)** Western blot of wildtype and motif-mutated, V5-tagged Tis11 from log-phase cells grown in complete synthetic media. **(B)** Growth curves for wildtype, motif mutant, and *Δtis11* in complete synthetic media alone (no ferrozine) and **(C)** with 750 µM ferrozine and 300 µM ammonium iron(II) sulfate. Different shades correspond to biological replicates (n=3) **(D)** Fragments from Eap1 in wildtype disordered library screen, as in Figure 4C. The Y(X_4_)Lφ and repressive motif regions are highlighted in purple. **(E)** Sequence alignment of conserved sequences within the repressive region of Eap1. Motif-1 and Motif-2 are displayed and correspond to naming in Figure 5G-H.

**Figure S6:**
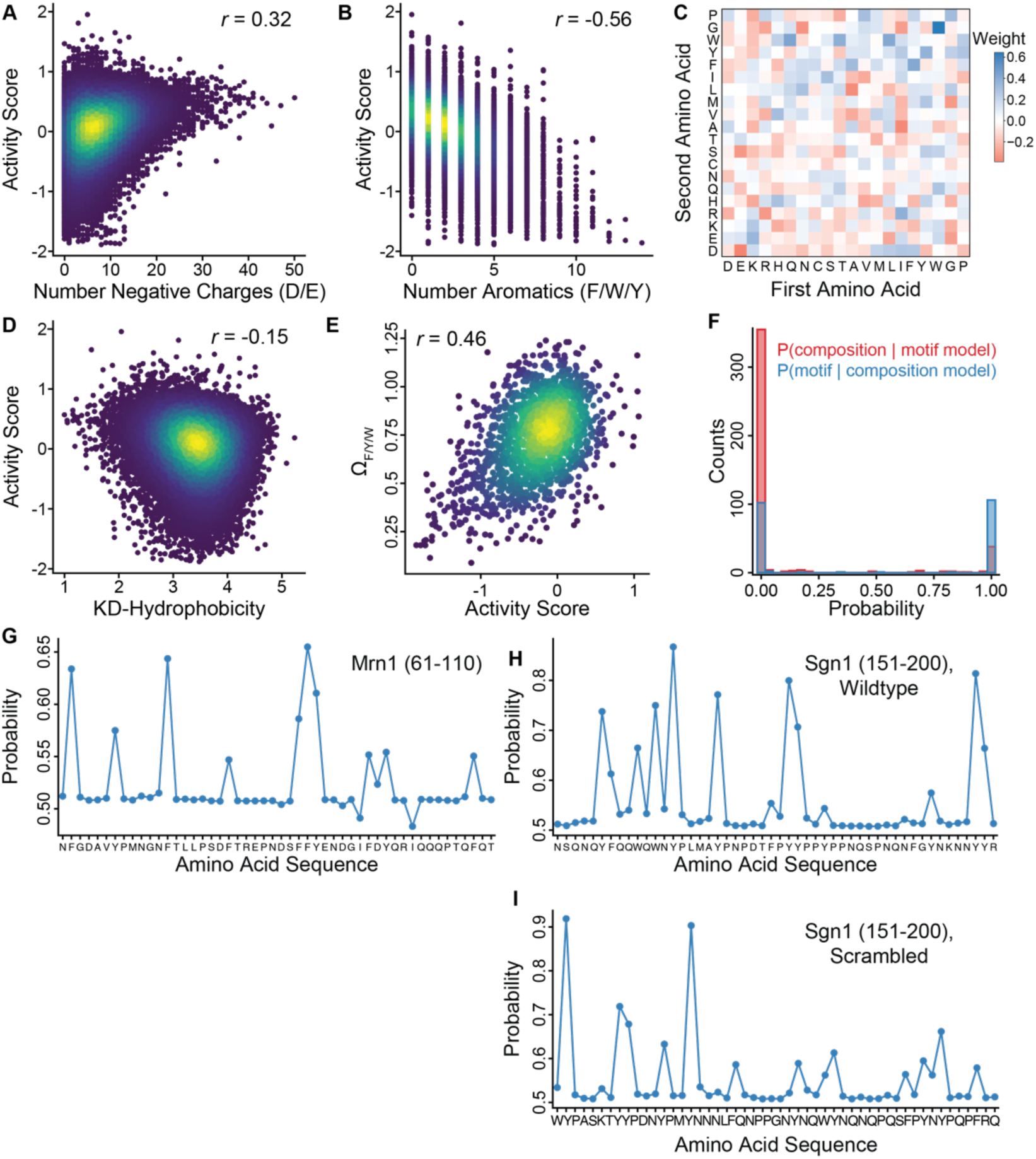
Features and predictions of repressive post-transcriptional disordered regions. **(A)** Correlation between number of negative changes and average activity scores (n=2). **(B)** Correlation between number of aromatic residues and average activity scores (n=2). **(C)** Coefficients for dipeptides in the logistic regression classifier for composition-driven repressors. The model includes both single-amino acid and dipeptides as variables. **(D)** Correlation between activity score and Kyte-Doolittle hydrophobicity. **(E)** Correlation between average activity scores for composition-driven repressors and the spacing of aromatic amino acids as defined by the parameter Ω_aro_ (n=2). Activity score is shown for each wildtype and stable scrambled composition-driven peptide, determined from the mutational scanning screen. **(F)** Prediction probabilities of the motif-dependent classifier to predict the composition-driven repressors (red) and the composition-driven classifier to predict motif-dependent repressors (blue). **(G-I)** Light attention model predictions for three composition-driven repressors using their ESM-1b representations. Peptide examined is displayed in panel legend.

## Materials and Methods

### Yeast strain construction

All *S.cerevisae* strains used were derived from BY4742 and transformed with the LiAc/PEG method^65^. Genomic integrations were verified by PCR amplification across sequence junctions. Unless otherwise stated, all yeast were grown at 30 °C. For construction of the reporter strain yJL004 containing the YFP-5xBoxB reporter and the mScarlet-5xPP7 normalizer, a codon optimized citrine containing 5x boxB hairpins, or mScarlet with 5x PP7 hairpins, were cloned into separate easyclone 2.0 vectors through Gibson assembly to make pJL001 and pJL002^66^. These vectors were linearized through NotI digestion for 1 hr at 37 °C before transforming log phase yeast with these constructs using the LiAc/PEG approach. To construct auxin inducible degron strains, *Os*TIR1 was cloned into an easyclone vector, creating pJL007, which was linearized by NotI digestion and transformed into yJL004 to create yJL009. CRISPR editing was used to introduce the mAID tag onto the C-terminus of Ccr4, Pop2, or Dcp2 at the endogenous locus. Guides targeting the C-terminus of the target protein were introduced into through golden gate cloning into pJR3428, which is a vector containing Cas9 and *URA3*, and repair templates containing synonymous mutations at the guide sites and the V5-mAID were ordered (Twist Biosciences), amplified by PCR, and co-transformed along with the pJL3428 expressing the appropriate guide into yJL009. Introduction of the mAID tag was verified by PCR, and the CRISPR plasmid was removed by patching positive colonies onto a YEPD plate followed by streaking onto 5-FOA plates. Tagged *tis11* strains were generated by similar methods through insertion of an N-terminal V5 tag and mutation of residues within the identified motif if needed. The knockout *tis11* was produced by replacing the *tis11* coding region with a hygromycin resistance cassette through homologous recombination. All yeast strains, plasmids, and oligos used in this study are available in Table S1.

### Flow cytometry

We used Gibson assembly to introduce a λN-emiRFP670 fusion expressed by the constitutive PGK1 promoter on a CEN/ARS plasmid to create pJL022, which was used in all tethering assays. Individual constructs were cloned into pJL022 by linearizing with BlpI and Gibson assembly of the appropriate peptide. The yJL004 tethering reporter stain was transformed with these constructs as described above^28^. At least two biological replicates from independent yeast colonies were chosen and grown overnight in SCD-Ura, back-diluted to OD_600_ = 0.05, and then grown to OD_600_ = 0.5-0.8. Yeast cells were pelleted by centrifugation at room temperature at 3000 g for 5 min, washed with PBS, and fixed in 4% paraformaldehyde/PBS solution at room temperature in the dark for 15-60 min. Fixed cells were pelleted, resuspended in fresh PBS, and stored at 4 °C prior to analysis by flow cytometry. Cells were transferred to polystyrene tubes and flow cytometry was performed measuring YFP, RFP, and iRFP fluorescence on ∼10K cells after gating on side and forward scatter, on both height and area, using a BD LSRFortessa cell analyzer. Flow cytometry data was analyzed with flowCore in R after applying the gating schemes on the forward and side scattering that selected for single cells^67^. We calculated the YFP/RFP ratio for each cell and reported the mean YFP/RFP or iRFP/RFP ratio for each replicate. The average of 2-3 biological replicates were reported, and sample sizes are indicated in the figure legend.

### Design and construction of disordered region library

For the initial library of peptides from all disordered regions, we downloaded all non-dubious ORFs from the Saccharomyces Genome Database and used IUPred2a to predict the disordered propensity of every residue within each gene^68^. We used a python workflow to tile across each ORF with a 50 amino acid window in one residue increments and removed windows with an average predicted disordered < 0.5. We eliminated windows less than 10 amino acids apart, except when the window spanned the last 50 amino acids of the ORF. This approach yielded a library of 46,473 unique sequences, which were reverse translated and codon optimized for yeast expression by Twist Biosciences. 24-25nt overhangs were added to these sequences to facilitate cloning, and oligos were synthesized as a pooled library for use in Gibson assembly. All sequences for the disordered region library are available in Table S2.

Library pools were amplified using KAPA HiFi Hotstart Readymix (Roche KK2601) with 0.3 µM each oJL109 and oJL090 and 20 ng input DNA per 50 µL PCR. Two 50 µL reactions were performed with 95 °C for 3 min, followed by 10-12 cycles of 98 °C for 20 s, 60 °C for 15 s, and 72 °C for 15 s, with a final extension of 1 min at 72 °C. The PCR was analyzed by gel electrophoresis and purified by spin column. To generate the library, 3 µg of pJL022 were digested with BlpI for 1 hr at 37 °C followed by spin column purification. 1 µg of linearized pJL022 and 144 ng of PCR amplified library were combined in 50 µL, mixed with 50 µL 2x NEBuilder HiFi DNA assembly Master mix (NEB E2621L) and incubated at 50 °C for 2 hr. The Gibson assembly was purified by spin column, eluted in 7 µL of water, and electroporated into 5x20 µL of MegaX DH10B T1^R^ Electrocomp cells (ThermoFisher C640003) according to the manufacturers protocol. After recovery at 37 °C for 1 hr, cells were transferred into 200 mL LB+Carb and grown overnight at 28 °C to OD_600_ = 3.0. Library coverage was estimated by serial dilution of transformations prior to overnight growth, plating dilutions onto LB+Carb, incubating overnight at 37 °C, and counting colonies to ensure the library contained >100 unique transformants for each oligo in the pool. The library was then purified from 200 mL overnight cultures by Midiprep (Qiagen 12143) according to manufacturer’s protocol.

### Pooled tethering assay and FACS

Reporter yeast strains were transformed with the appropriate library by growing 250 mL of yeast in YEPD to OD_600_ = 0.5. Specifically, cells were collected by centrifugation and transformed using a scaled up LiAc/PEG procedure with >30 µg of library DNA for each of two biological replicates. The yeast were grown for two days in 500 mL SCD-Ura at 22 °C until OD_600_ = 1.5-3, and 210 OD units were collected by centrifugation at 3000 g for 5 min at room temperature and stored at -80 °C in SCD-Ura+15% glycerol until ready for use. Coverage was estimated by serial dilution of transformations prior to selective outgrowth, plated on SC-Ura, and incubated 2 days at 30 °C to ensure that coverage of >10x of each oligo in the library.

To perform screens, glycerol stocks were thawed, collected by centrifugation at 3000 g for 5 min, and washed three times with 10 mL SCD-Ura. Cells were transferred into a custom turbidostat and grown at constant turbidity to achieve steady state levels of YFP/RFP fluorescence and remove non-transformed yeast from the culture^31^. Cells were grown for ≥16 hr at 30 °C in log-phase (OD_600_ = ∼1) with constant aeration and agitation. 50 mL of cells were harvested by centrifugation at 3000 g for 5 min and washed once with cold PBS before being resuspended in ∼20 mL cold PBS. Cells were then sorted on a BD FACSAria using the gating schemes described above. For both the YFP/RFP and iRFP/RFP sorting, populations were divided into four gates with equal number of cells and > 1 million cells were collected in each bin. 2 mL YEPD was added to the sorted cells, as well as two unsorted populations for each replicate, and transferred to RNAse-free tubes before collecting cells by centrifugation at 3000 g for 5 min. The supernatant was removed and cells were flash frozen in LN_2_ and stored at -80 °C prior to RNA extraction. Tethering assays conducted in degron-tagged yeast strains were performed as described above, but with SCD-Ura supplemented with indole-3-acetic acid (IAA, Sigma I2886) to 0.5 mM.

### RNA extraction, library preparation, and DNA sequencing

Each peptide is encoded by a unique nucleotide sequence that can be identified by short read sequencing. We chose to isolate mRNA and prepare libraries from cDNA rather than extract plasmids, as this approach avoids bottlenecks associated with low plasmid recovery rates^69^. To isolate mRNA, cell pellets were thawed and RNA was extracted using the hot phenol approach^70^. Specifically, cells pellets were resuspended in 440 µL of 50 mM NaOAc pH 5.2, 10 mM EDTA, 1% SDS (v/v) and 400 µL phenol/chloroform. The mixture was heated with agitation at 65 °C for 15 minutes before being cooled on ice for 5 min. The aqueous phase was separated by centrifugation at 20,000 g at 4 °C for 5 min and re-extracted twice with 400 µL chloroform, spinning for 5 minutes at 20,000 g after each extraction. The aqueous phase was transferred to a new RNAse-free tube and RNA was precipitated by adding 50 µL of 3 M NaOAc, pH 5.2, 1 µL of GlycolBlue (Invitrogen, AM9515), and EtOH to 80% (v/v) to the sample before chilling at -80 °C for 30 minutes. The RNA pellet was collected by centrifugation at 20,000 g, 4 °C for 10 min and washed once with 500 µL ice-cold 70% EtOH. The pellet was then air dried for 5 min and resuspend in 40 µL water. The RNA was treated with TURBO DNAse (Invitrogen, AM2238) in a 50 µL final volume for 30 min at 37 °C. The sample was then adjusted to 180 µL with water, followed by addition of 20 µL 3 M NaOAc, pH 5.2 and 200 µL of phenol/chloroform. The aqueous phase was separated by centrifugation at 20,000 g for 5 min, re-extracted twice with 200 µL with chloroform, and precipitated with ethanol as described above. The RNA pellet was resuspended in 20 µL of water and quantified by NanoDrop spectrophotometry. Reverse transcription was performed with dT priming using the ProtoScript II RT (NEB M0368L) as described by the manufacturer, scaled up two-fold. RNA was the digested by adding 0.5 µL each of RNAse A and RNAse H (ThermoFisher R1253 and NEB M0297L) to each reaction and incubating for 30 min at 37 °C, followed by spin column purification (Qiagen, 28104). The cDNA was then used as input for the first-round PCR with oJL105 and oJL106 to add the i5/i7 primer sites using Q5 DNA polymerase (NEB M0491L). Libraries were amplified with an initial 98 °C denaturation step for 30 s, followed by 8-10 cycles of 98 °C for 10 s, 60 °C for 20 s, and 72 °C for 10 s, with a final extension time for 2 min. The first-round product was purified by spin column, and half the elution was used as a template in a second round of 8-10 cycle PCR to prepare Illumina libraries using NEBNext multiplex oligos using the same conditions described above (NEB E7335L and E7500L). The final PCR product was purified using AMPure XP beads (Beckman Coulter A63882) at a 1:1 ratio with the PCR product according to the manufacturers protocol and resuspend in 25 µL water. PCR products were verified by automated electrophoresis using an Agilent Tapestation 2200 and libraries were pooled at equimolar ratio. For with proteome-wide disordered peptide library, sequencing was performed on an Illumina HiSeq 4000 instrument using single-end reads. For the mutational scanning library, sequencing was performed on an Illumina HiSeq 4000 with paired-end reads. We targeted ∼5 million reads for each gated population with the sorted and unsorted samples.

### Deep sequencing processing and determination of activity and stability scores

Illumina sequencing reads were trimmed with cutadapt, mapped to the oligo library using bowtie2 and counts were extracted from alignments with samtools^71–73^. Oligos with fewer than 25 reads in the unsorted sample, or less than 5 total reads in the sorted samples, were discarded and excluded from scoring. Reads in each bin were normalized to the total number of reads in the sample, and then activity scores were determined through a maximum likelihood estimator using a custom R script and scoring function based on established procedures^29^. Stability scores were determined exactly like activity scores, except using the iRFP/RFP sorted samples. Fragments that had an activity score ≤ -1 were included, regardless of stability score, as these had repressive activity and were therefore stable enough to induce their regulatory effects. For fragments with activity scores > -1, we categorized fragments as unstable if they had a stability score ≤ -1. These fragments could be truly nonfunctional or simply expressed to poorly to observe activity. For gene-level plots, fragments from a protein were grouped and colored conditionally based on their stability scores.

### Gene ontology and RBP enrichment analysis

We determined fragments with activity scores ≤ -1 and categorized these as repressors. We then took the full list of proteins that had at least one peptide present in our analysis and used these as the background gene list. These sets were imported into PantherDB to calculate statistical overrepresentation using the Fisher’s exact test corrected by the calculated false discovery rate^74^. We curated select GO terms with ≥ 3.5-fold enrichment to display and included the full list of terms as supplemental data. Enrichment of yeast RBPs was determined by comparing proteins with at least one repressive fragment to a collection of different yeast-RBP mass spectrometry datasets^9^. We compared the activity scores to the number of different studies that experimentally identified a protein as an RBP. We reasoned that the presence of a particular protein in multiple studies provided higher confidence of a true RBP.

### Degron tagging

A C-terminal V5-mAID tag was added to the endogenous Pop2, Ccr4, or Dcp2 in yJL009 through CRISPR-based gene editing as described above to create yJL026, yJL027, and yJL010, respectively. Fitness effects were measured in biological triplicate by growing cells in YEPD overnight before diluting into YEPD at OD_600_ = 0.1 and growing for 2-3 doublings. Cells were then again diluted to OD_600_ = 0.05 in YEPD with either 1% DMSO or 0.5 mM IAA final concentration and 100 µL was transferred to a 96 well U-bottom plate before covering with Breathe-Easy gas permeable seal. Cells were grown at 30 °C with constant agitation in a Tecan Spark plate reader while measuring the OD_600_ every 5 minutes. Growth curves were fit to the logistic growth equation:

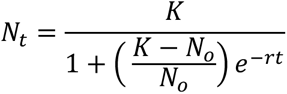

Where *K, N_o_, N_t_, r,* and *t* are the maximum OD_600_, the initial OD_600_, the OD_600_ at time *t*, the growth rate, and the time, respectively. The parameters *K, N_o_* and *r* were fit simultaneously using custom scripts written in R.

Depletion of the mAID tagged protein was verified by western blotting. Cells were grown overnight in YEPD, diluted to OD_600_ = 0.05, and grown for 2-3 doublings, after which IAA (0.5 mM final concentration) or DMSO (1%, v/v) was added to culture before growing for another 4 hours. Yeast were collected by centrifugation at 3000 g for 5 min, resuspended in 500 µL 20% trichloroacetic acid (TCA), and pelleted by centrifugation at 16,000 g for 2 min. The TCA was removed and pellets were flash frozen in LN_2_ and stored at -80 °C until ready to use. Pellets were then thawed on ice for 5 min before adding 250 µL 20 % TCA and 250 µL 0.5 mm glass beads. The samples were vortexed 3x1 min at room temperature, with chilling on ice for 1 minute between cycles of bead beating. Beads were clarified by the double-tube method and then washed once 300 µL 5 % TCA. After removing beads, 700 µL of 5% TCA was added to the sample and protein was pelleted by spinning at 12,000 g for 10 min at 4 °C. The TCA was then removed and the pellet was washed with 750 µL ice-cold 100% ethanol before another 10 min centrifugation at 12,000 g at 4 °C. The ethanol was discarded, and the pellet was allowed to air dry for 5 min before resuspending in 0.1 M Tris pH 8.0 and stored at -20 °C until ready to use. Protein was quantified by A_280_ absorbance, and 80 µg of protein was loaded onto a 4-12% Bis-Tris polyacrylamide gel and separated by electrophoresis before being transferred to a 0.2 µm PVDF membrane. The membrane was blocked with 5% milk and probed using either 1:1000 antiV5 antibody (Cell Signaling Technology, 13202) or 1:10000 GAPDH as a loading control (Proteintech 60004-1). Blots were developed with SuperSignal West Dura Extended Duration Substrate (Thermo Scientific 34075) and luminescence was imaged using a ProteinSimple (Bio-techne).

### Epistasis analysis

Growth of degron tagged yeast, FACS, and library preparation was performed as described above in two biological replicates. For degron tagged yeast grown in 0.5 mM IAA, cells were only sorted on the YFP/RFP ratio. We determined the activity scores for each peptide using our maximum likelihood estimation approach as described above and excluded peptides that had a stability score ≤ -1 in the wildtype background. The mean activity scores of the degron tagged strains were shifted relative to the wildtype, which can occur due to fitness defects and/or differences in the underlying fluorescence distribution. To account for this, we assume that the true activity of most peptides does not change in the mutant relative to the wildtype and thus rescaled the activity scores of peptides in each replicate of the degron-tagged stain based on LOESS regression against the average wildtype activity scores. We then determined statistically different activities using the Z-test and adjusted p-values using the Benjamini-Hochberg procedure. For comparing between degron strains, we only included peptides with sufficient coverage in the wildtype strain and all three degron tagged mutants. The UpSetR package was used to visualize clusters of peptides that had differential activity between the wildtype and degron-tagged background^75^.

### Mutational scanning library generation

For the mutational scanning library, we selected all peptides that had an activity score ≤ -1 in our high-throughput tethering assay as candidates. If two peptides overlapped by > 30 residues, the more repressive of the two fragments was included. This produced a library of 699 fragments for mutational scanning, including 20 inactivate controls (activity scores > -1). We replaced consecutive 5 amino acid blocks with a GGSSG sliding window in one amino acid increments, as well as included 5 randomly shuffled sequences. We also included fragments from orthologs from known yeast species, when present, that aligned with these repressors. While these fragments were included in our library, we did not further analyze them and they were only used to calculate a population standard deviation for subsequent Z-test scores. The shuffled sequences and orthologs were reverse translated and codon optimized by Twist Biosciences, while the 15-nucleotide constant sequence encoding GGSSG substitution was tiled across the original repressor sequence. The mutational scanning library was constructed exactly as described for the disordered yeast proteome library. The reporter yeast strain, yJL004, was then transformed with this library followed by sorting cells and scoring activity and stability as described above. All sequences for the mutational scanning library are available in Table S3.

### Mutational scanning with Hidden Markov Models and extraction of functional motifs

Activity scores of variants produced by mutational scanning were fit to hidden Markov models (HMMs) using depmixS4 in R^76^. Mutational scanning variants for each repressor were grouped by the parent peptide, and fragments with stability scores ≤ -1 were excluded. Mutational scanning profiles were then fit to a two state HMM using the Viterbi algorithm. Three repressors that failed to converge during fitting were excluded from analysis. Mutational windows associated with the states with a more negative activity score were assigned as inactive regions, as these windows did not affect activity of the repressor. Conversely, mutational regions with a higher activity score (less repressive) corresponded to active regions because mutating these residues abolished repressor function. Motif-dependent HMMs were defined by repressors that had statistically significant loss of activity in the scrambled variants (see below).

Functional subregions were extracted by isolating continuous regions that were disrupted by the GGSSG mutation. We only examined fragments that had fewer than 5 transitions within the HMM and had at least two consecutive mutational windows in both the active and inactive state. Because we used a 5 amino acid GGSSG mutational scanning, we only included residues with the active regions that started at *i+*4, where *i* is the start of the mutational window within the repressor, and ended at the last residue within this region. We assumed all residues from 46-50 belonged to the same state as the last mutation window and excluded subregions shorter than 3 amino acids for downstream analysis.

### Amino acid enrichment analysis

Amino acid enrichment analysis was performed using Composition Profiler with default settings for bootstrap iterations (10,000) and significance cutoff (0.05)^77^. For computing the enrichment of amino acids in the motifs to background, we compared the residues in mutationally sensitive regions to the amino acids in the mutationally tolerant windows. The full disordered library was highly overlapping and could lead to overcounting residues. To eliminate this problem, we reconstructed the contagious disordered regions for each ORF from the overlapping fragments. We then compared the enrichment of these disordered regions to the non-dubious ORFs from the Saccharomyces genome database.

### Motif visualization, alignment, and structural modelling

The Tis11 and Eap1 motifs were aligned using MUSCLE with yeast sequences from the yeast genome order browser and visualized using Jalview^78–80^. For the Edc1 motif, the homologous human PNRC2 motif was used as a query in MotifFinder and selected sequences were used for alignment. Motifs within Eap1 were identified by sequence alignment with related yeast species.

For AlphaFold2 modelling of the Tis11/Cdc39 (Not1) interaction, residues 587-758 of Cdc39 and residues 28-59 of Tis11 were used with default settings using ColabFold^81^. The best model was selected manually and displayed using PyMol.

### Iron depletion and growth assays with Tis11 strains

Growth assays were performed in biological triplicates, and strains were grown overnight in at 30 °C in SCD-complete media and diluted to OD_600_ = 0.1 in the appropriate media. For conditions with ferrozine or supplemental iron, SCD-complete was supplemented with 750 µM Ferrozine +/- 300 µM Ammonium iron(II) sulfate (Sigma 160601 and 215406). Cells were grown at 30 °C until OD_600_ = ∼ 0.5 and further diluted to OD_600_ = 0.05 in the appropriate media. Cells were then grown in a Tecan Spark plate reader as described above for 24 hr at 30 °C with constant agitation while recording the OD_600_ every 5 minutes. For western blot of wildtype and mutant Tis11, yeast were harvested from 5 mL of log-phase culture and protein was isolated by TCA precipitation and blotted as described above for both V5 and Hxk2 as a loading control (Rockland #100-4159).

### Identifying compositional-dependent repressors

We assumed that compositional-dependent repressors should retain activity when the sequence was shuffled. Therefore, we determined the average score of the scrambles of each wildtype repressor sequence. After discarding repressors with < 2 stable scrambles (stability scores ≤ -1), we then performed a Z-test between the average activity score of these scrambled fragments and the activity score of the wildtype repressor. We chose to use the activity scores determined by the HMM analysis fit to all the mutant fragments, as this was more statistically robust than the single wildtype repressor within our screen. Fragments where the activity scores did not differ significantly between the scramble and the wildtype sequence (Benjamini-Hochberg corrected p-values > 0.05) were identified as composition-dependent fragments. Eleven repressors had average scramble scores that were statistically more repressive than the wildtype peptide, which were excluded from downstream analysis.

### Machine learning

Logistic regression analysis on the amino acid composition of repressors was performed using the *scikit-learn* package in Python^82^. Repressors were assigned as the positive class (1), and the inactive fragments from the wildtype disordered library screen were the negative class (0). We excluded unstable inactive fragments (stability score ≤ -1) and fragments with an activity score greater than -1 and less that -0.75 during classification. We fit the composition-dependent peptides and repressive motifs separately; for the motifs, we only used the active subregion. We removed inactive fragments that overlapped with repressors and fragments that overlapped by > 30 amino acids. We then performed logistic regression using either the primary amino acid sequence (20 parameters) or included dipeptides (420 total parameters). For fitting to the primary amino acid sequence, we used the fraction of each 20 amino acid in the sequence, which allowed us to compare models where repressors had different sequence lengths. We only performed dipeptide analysis for the composition dependent fragments, as there were too few dipeptides in the motifs for analysis. In the dipeptide analysis, we used the number of occurrences of each single and dipeptide in the model.

To perform the logistic regression analysis, data was split using an 80/20 train/test set and fit to logistic model using *liblinear* solver with L2 regularization and 5-fold cross validation. The L2 penalty, *λ*, and class weights were selected from a grid search of 40 and 20 logarithmically spaced starting values, respectively, and evaluated based on area under the precision recall curve (AUPRC). The best parameters were then used in the final model, and the precision, recall and AUPRC were exported for plotting in R.

The light attention model was performed using PyTorch on the ESM-1b embeddings of the compositional dependent repressors. Each sequence was represented as a 50 x 1280 embedding with the *esm1b_t33_650M_UR50s* pre-trained model using the *esm-extract* command with the *per_tok* setting^54^. The final layer of the embedding was loaded and used as inputs for a light attention PyTorch model. Because these embeddings are sensitive to the position of each amino acid, the composition-dependent wildtype fragments and their associated scrambles have different representations, unlike the primary amino acid sequence, which is the same between the wildtype and the shuffled sequence. To avoid potential memorization, the scrambles and wildtype from the same parent fragment were always grouped together in the same training or test set, using an 80/20 split. The inactive controls were the same as used in the logistic regression model.

This model performs the following operations to weight each row of any N x 1280 representation, where N is the number of residues of the protein sequence used to generate the representation. First, this model computes the dot product between a 1 x 1280 trainable weighting vector and each row of the embedding to produce a N x 1 weighting vector, which is then normalized through a Softmax function. Each element in row *i* of the N x 1280 embedding is multiplied by the *i^th^* element of the Softmax-normalized weighting vector to weight the rows. After weighting, the N x 1280 embedding is summed column-wise to produce a final 1 x 1280 vector, which is then multiplied by a learned linear layer before being passed through a sigmoid activation function to produce a final classification.

Both the learning rate and L2 penalty were grid searched from 10 logarithmically spaced values and each model was trained for 2000 epochs using the Adam optimizer to minimize a binary cross entropy loss function. The learning rate and L2 penalty producing the best AUPRC were chosen for the final model, which was trained for 5000 epochs. For predicting individual position weights within each amino acid, these weights were loaded into a model that worked essentially as described above, but applied the linear weights and sigmoid activation function to each row of an N x 1280 representation to produce an N x 1 prediction vector, where each element is the probability of that residue having repressive activity.

### Physicochemical property calculations

The total counts for acidic or aromatic amino acids were the sum of the number of D/E or Y/F/W residues in each fragment. The Kyte-Doolittle hydrophobicity was calculated using the *localCIDER* package in Python^83^. The Ω_aro_ for each repressor was calculated as described previously, using a blob length of 5 residues^53^.

### Quantification and statistical analysis

Statistical analyses were performed in Python 3.8.13 and R-4.2.0. Two-sided Z-tests were used when more than 30 data points were sampled. We used a Benjamini-Hochberg corrected p value of ≤ 0.05 to assess significance when appropriate.

### Data and code availability

Raw sequencing data and processed reads generated from this study has been deposited in the NCBI Gene Expression Omnibus (accession number GSE254492). Code, analysis, and raw data is available on Github (https://github.com/ingolia-lab/post-transcriptional-idrs).

## References

1. Wright, P. E. & Dyson, H. J. Intrinsically disordered proteins in cellular signalling and regulation. Nat. Rev. Mol. Cell Biol. 16, 18–29 (2015).

2. Holehouse, A. S. & Kragelund, B. B. The molecular basis for cellular function of intrinsically disordered protein regions. Nat. Rev. Mol. Cell Biol. (2023).

3. Staller, M. V. et al. A High-Throughput Mutational Scan of an Intrinsically Disordered Acidic Transcriptional Activation Domain. Cell Syst. 6, 444–455 (2018).

4. Erijman, A. et al. A High-Throughput Screen for Transcription Activation Domains Reveals Their Sequence Features and Permits Prediction by Deep Learning. Mol. Cell 78, 890–902 (2020).

5. Sanborn, A. L. et al. Simple biochemical features underlie transcriptional activation domain diversity and dynamic, fuzzy binding to Mediator. Elife 10, 1–42 (2021).

6. Van Roey, K. et al. Short linear motifs: Ubiquitous and functionally diverse protein interaction modules directing cell regulation. Chem. Rev. 114, 6733–6778 (2014).

7. Calabretta, S. & Richard, S. Emerging Roles of Disordered Sequences in RNA-Binding Proteins. Trends Biochem. Sci. 40, 662–672 (2015).

8. Jonas, S. & Izaurralde, E. The role of disordered protein regions in the assembly of decapping complexes and RNP granules. Genes Dev. 27, 2628–2641 (2013).

9. Hentze, M. W., Castello, A., Schwarzl, T. & Preiss, T. A brave new world of RNA-binding proteins. Nat. Rev. Mol. Cell Biol. 19, 327–341 (2018).

10. Calabretta, S. & Richard, S. Emerging Roles of Disordered Sequences in RNA-Binding Proteins. Trends Biochem. Sci. 40, 662–672 (2015).

11. Brent, R. & Ptashne, M. A eukaryotic transcriptional activator bearing the DNA specificity of a prokaryotic repressor. Cell 43, 729–736 (1985).

12. Reynaud, K., McGeachy, A., Noble, D., Meacham, Z. & Ingolia, N. Surveying the global landscape of post-transcriptional regulators. Nat. Struct. Mol. Biol. (2023).

13. Luo, E. C. et al. Large-scale tethered function assays identify factors that regulate mRNA stability and translation. Nat. Struct. Mol. Biol. 27, 989–1000 (2020).

14. Erben, E. D., Fadda, A., Lueong, S., Hoheisel, J. D. & Clayton, C. A Genome-Wide Tethering Screen Reveals Novel Potential Post-Transcriptional Regulators in Trypanosoma brucei. PLoS Pathog. 10, (2014).

15. Van Nostrand, E. L. et al. A large-scale binding and functional map of human RNA-binding proteins. Nature 583, 711–719 (2020).

16. Dominguez, D. et al. Sequence, Structure, and Context Preferences of Human RNA Binding Proteins. Mol. Cell 70, 854–867 (2018).

17. Zarin, T. et al. Proteome-wide signatures of function in highly diverged intrinsically disordered regions. Elife 8, 1–26 (2019).

18. Raisch, T. et al. Distinct modes of recruitment of the CCR 4– NOT complex by Drosophila and vertebrate Nanos. EMBO J. 35, 974–990 (2016).

19. Peter, D. et al. Molecular Architecture of 4E-BP Translational Inhibitors Bound to eIF4E. Mol. Cell 57, 1074–1087 (2015).

20. Schwartz, D., Decker, C. J. & Parker, R. The enhancer of decapping proteins, Edc1p and Edc2p, bind RNA and stimulate the activity of the decapping enzyme. RNA 9, 239–251 (2003).

21. Coller, J. M., Gray, N. K. & Wickens, M. P. mRNA stabilization by poly(A) binding protein is independent of poly(A) and requires translation. Genes Dev. 12, 3226–3235 (1998).

22. Koren, I. et al. The Eukaryotic Proteome Is Shaped by E3 Ubiquitin Ligases Targeting C-Terminal Degrons. Cell 173, 1622–1635 (2018).

23. Keryer-Bibens, C., Barreau, C. & Osborne, H. B. Tethering of proteins to RNAs by bacteriophage proteins. Biol. Cell 100, 125–138 (2008).

24. Bindels, D. S. et al. MScarlet: A bright monomeric red fluorescent protein for cellular imaging. Nat. Methods 14, 53–56 (2016).

25. DelRosso, N. et al. Large-scale mapping and mutagenesis of human transcriptional effector domains. Nature 616, 365–372 (2023).

26. Sheff, M. A. & Thorn, K. S. Optimized cassettes for fluorescent protein tagging in Saccharomyces cerevisiae. Yeast 21, 661–670 (2004).

27. Luke, B. et al. Saccharomyces cerevisiae Ebs1p is a putative ortholog of human Smg7 and promotes nonsense-mediated mRNA decay. Nucleic Acids Res. 35, 7688–7697 (2007).

28. Matlashov, M. E. et al. A set of monomeric near-infrared fluorescent proteins for multicolor imaging across scales. Nat. Commun. 11, 1–12 (2020).

29. Peterman, N. & Levine, E. Sort-seq under the hood: Implications of design choices on large-scale characterization of sequence-function relations. BMC Genomics 17, 1–17 (2016).

30. Dvir, S. et al. Deciphering the rules by which 5’-UTR sequences affect protein expression in yeast. Proc. Natl. Acad. Sci. U. S. A. 110, (2013).

31. McGeachy, A. M., Meacham, Z. A. & Ingolia, N. T. An Accessible Continuous-Culture Turbidostat for Pooled Analysis of Complex Libraries. ACS Synth. Biol. 8, 844–856 (2019).

32. Goldstrohm, A. C., Hook, B. A., Seay, D. J. & Wickens, M. PUF proteins bind Pop2p to regulate messenger RNAs. Nat. Struct. Mol. Biol. 13, 533–539 (2006).

33. Kilchert, C., Wittmann, S. & Vasiljeva, L. The regulation and functions of the nuclear RNA exosome complex. Nat. Rev. Mol. Cell Biol. 17, 227–239 (2016).

34. Bresson, S., Tuck, A., Staneva, D. & Tollervey, D. Nuclear RNA Decay Pathways Aid Rapid Remodeling of Gene Expression in Yeast. Mol. Cell 65, 787–800.e5 (2017).

35. Webster, M. W., Stowell, J. A. & Passmore, L. A. RNA-binding proteins distinguish between similar sequence motifs to promote targeted deadenylation by Ccr4-Not. Elife 8, 1–26 (2019).

36. Poetz, F. et al. RNF219 attenuates global mRNA decay through inhibition of CCR4-NOT complex-mediated deadenylation. 12, (2021).

37. Fabian, M. R. et al. Structural basis for the recruitment of the human CCR4-NOT deadenylase complex by tristetraprolin. Nat. Struct. Mol. Biol. 20, 735–739 (2013).

38. Keskeny, C. et al. A conserved CAF40-binding motif in metazoan NOT4 mediates association with the CCR4–NOT complex. Genes Dev. 33, 236–252 (2019).

39. Bhandari, D., Raisch, T., Weichenrieder, O., Jonas, S. & Izaurralde, E. Structural basis for the Nanos-mediated recruitment of the CCR4-NOT complex and translational repression. Genes Dev. 28, 888–901 (2014).

40. Parker, R. RNA degradation in Saccharomyces cerevisae. Genetics 191, 671–702 (2012).

41. Nishimura, K. & Kanemaki, M. T. Rapid depletion of budding yeast proteins via the fusion of an auxin-inducible degron (AID). Curr. Protoc. Cell Biol. 2014, 20.9.1–20.9.16 (2014).

42. Basquin, J. et al. Architecture of the nuclease module of the yeast ccr4-Not complex: The not1-caf1-ccr4 interaction. Mol. Cell 48, 207–218 (2012).

43. Wurm, J. P., Holdermann, I., Overbeck, J. H., Mayer, P. H. O. & Sprangers, R. Changes in conformational equilibria regulate the activity of the Dcp2 decapping enzyme. Proc. Natl. Acad. Sci. 114, 6034–6039 (2017).

44. Mugridge, J. S., Tibble, R. W., Ziemniak, M., Jemielity, J. & Gross, J. D. Structure of the activated Edc1-Dcp1-Dcp2-Edc3 mRNA decapping complex with substrate analog poised for catalysis. Nat. Commun. 9, (2018).

45. Varier, R. A. et al. m6A reader Pho92 is recruited co-transcriptionally and couples translation efficacy to mRNA decay to promote meiotic fitness in yeast. Elife 11, (2022).

46. Gruner, S. et al. Structural motifs in eIF4G and 4E-BPs modulate their binding to eIF4E to regulate translation initiation in yeast. Nucleic Acids Res. 46, 6893–6908 (2018).

47. Puig, S., Askeland, E. & Thiele, D. J. Coordinated remodeling of cellular metabolism during iron deficiency through targeted mRNA degradation. Cell 120, 99–110 (2005).

48. Prouteau, M., Daugeron, M. C. & Séraphin, B. Regulation of ARE transcript 3′ end processing by the yeast Cth2 mRNA decay factor. EMBO J. 27, 2966–2976 (2008).

49. Cosentino, G. P. et al. Eap1p, a Novel Eukaryotic Translation Initiation Factor 4E-Associated Protein in Saccharomyces cerevisiae. Mol. Cell. Biol. 20, 4604–4613 (2000).

50. Mader, S., Lee, H., Pause, A. & Sonenberg, N. The Translation Initiation Factor eIF-4E Binds to a Common Motif Shared by the Translation Factor eIF-4γ and the Translational Repressors 4E-Binding Proteins. Mol. Cell. Biol. 15, 4990–4997 (1995).

51. Blewett, N. H. & Goldstrohm, A. C. A Eukaryotic Translation Initiation Factor 4E-Binding Protein Promotes mRNA Decapping and Is Required for PUF Repression. Mol. Cell. Biol. 32, 4181–4194 (2012).

52. Cridge, A. G. et al. Identifying eIF4E-binding protein translationally-controlled transcripts reveals links to mRNAs bound by specific PUF proteins. Nucleic Acids Res. 38, 8039– 8050 (2010).

53. Martin, E. W. et al. Valence and patterning of aromatic residues determine the phase behavior of prion-like domains. Science (80-.). 367, 694–699 (2020).

54. Rives, A. et al. Biological structure and function emerge from scaling unsupervised learning to 250 million protein sequences. Proc. Natl. Acad. Sci. U. S. A. 118, (2021).

55. Meier, J. et al. Language models enable zero-shot prediction of the effects of mutations on protein function. Adv. Neural Inf. Process. Syst. 35, 29287–29303 (2021).

56. Stärk, H., Dallago, C., Heinzinger, M. & Rost, B. Light attention predicts protein location from the language of life. Bioinforma. Adv. 1, 1–8 (2021).

57. Tycko, J. et al. High-Throughput Discovery and Characterization of Human Transcriptional Effectors. Cell 183, 2020–2035 (2020).

58. Arnold, C. D. et al. A high-throughput method to identify trans-activation domains within transcription factor sequences. EMBO J. 37, 1–13 (2018).

59. Ravarani, C. N. et al. High-throughput discovery of functional disordered regions: investigation of transactivation domains. Mol. Syst. Biol. 14, 1–14 (2018).

60. Tuttle, L. M. et al. Mediator subunit Med15 dictates the conserved “fuzzy” binding mechanism of yeast transcription activators Gal4 and Gcn4. Nat. Commun. 12, (2021).

61. Sharma, R., Raduly, Z., Miskei, M. & Fuxreiter, M. Fuzzy complexes: Specific binding without complete folding. FEBS Lett. 589, 2533–2542 (2015).

62. Zarin, T. et al. Identifying molecular features that are associated with biological function of intrinsically disordered protein regions. Elife 10, 1–23 (2020).

63. Vacic, V. et al. Disease-Associated Mutations Disrupt Functionally Important Regions of Intrinsic Protein Disorder. PLoS Comput. Biol. 8, (2012).

64. Mashahreh, B. et al. Conserved degronome features governing quality control associated proteolysis. Nat. Commun. 13, (2022).

65. Gietz, R. D. & Schiestl, R. H. High-efficiency yeast transformation using the LiAc/SS carrier DNA/PEG method. Nat. Protoc. 2, 31–34 (2007).

66. Stovicek, V., Borja, G. M., Forster, J. & Borodina, I. EasyClone 2.0: expanded toolkit of integrative vectors for stable gene expression in industrial Saccharomyces cerevisiae strains. J. Ind. Microbiol. Biotechnol. 42, 1519–1531 (2015).

67. Hahne, F. et al. flowCore: A Bioconductor package for high throughput flow cytometry. BMC Bioinformatics 10, (2009).

68. Mészáros, B., Erdös, G. & Dosztányi, Z. IUPred2A: Context-dependent prediction of protein disorder as a function of redox state and protein binding. Nucleic Acids Res. 46, W329–W337 (2018).

69. Muller, R., Meacham, Z., Ferguson, L. & Ingolia, N. CiBER-seq dissects genetic networks by quantitative CRISPRi profiling of expression phenotypes. Science (80-.). 370, (2020).

70. Green, M. R. & Sambrook, J. Total RNA Extraction from Saccharomyces cerevisiae using hot acid phenol. Cold Spring Harb. Protoc. 2021, 523–525 (2021).

71. Li, H. et al. The Sequence Alignment/Map format and SAMtools. Bioinformatics 25, 2078–2079 (2009).

72. Langmead, B. & Salzberg, S. L. Fast gapped-read alignment with Bowtie 2. Nat. Methods 9, 357–359 (2012).

73. Martin, M. Cutadapt removes adapter sequences from high-throughput sequencing reads. EMBnet.journal 17, (2011).

74. Thomas, P. D. et al. PANTHER: Making genome-scale phylogenetics accessible to all. Protein Sci. 31, 8–22 (2022).

75. Conway, J. R., Lex, A. & Gehlenborg, N. UpSetR: An R package for the visualization of intersecting sets and their properties. Bioinformatics 33, 2938–2940 (2017).

76. Visser, I. & Speekenbrink, M. depmixS4: An R package for hidden markov models. J. Stat. Softw. 36, 1–21 (2010).

77. Vacic, V., Uversky, V. N., Dunker, A. K. & Lonardi, S. Composition Profiler: A tool for discovery and visualization of amino acid composition differences. BMC Bioinformatics 8, 1–7 (2007).

78. Byrne, K. P. & Wolfe, K. H. The Yeast Gene Order Browser: Combining curated homology and syntenic context reveals gene fate in polyploid species. Genome Res. 15, 1456–1461 (2005).

79. Waterhouse, A. M., Procter, J. B., Martin, D. M. A., Clamp, M. & Barton, G. J. Jalview Version 2-A multiple sequence alignment editor and analysis workbench. Bioinformatics 25, 1189–1191 (2009).

80. Edgar, R. C. MUSCLE: Multiple sequence alignment with high accuracy and high throughput. Nucleic Acids Res. 32, 1792–1797 (2004).

81. Mirdita, M. et al. ColabFold: making protein folding accessible to all. Nat. Methods 19, 679–682 (2022).

82. Barupal, D. K. & Fiehn, O. Scikit-learn: Machine Learning in Python. J. Mach. Learn. Res. 12, 2825–2830 (2011).

83. Holehouse, A. S., Das, R. K., Ahad, J. N., Richardson, M. O. G. & Pappu, R. V. CIDER: Resources to Analyze Sequence-Ensemble Relationships of Intrinsically Disordered Proteins. Biophys. J. 112, 16–21 (2017).

